# Somatic mutation inference from single-cell transcriptomics: A survey in the esophagus

**DOI:** 10.64898/2026.07.07.737010

**Authors:** A. Méndez-Alejandre, D. González-Menéndez, S. Vidal-Notari, G. Skrupskelyte, M. Rodríguez-Rodríguez, H. Ajith, A. S. Torralba, M. P. Alcolea, G. Piedrafita

## Abstract

Human somatic tissues accumulate mutations during normal aging. Some of these affect cancer-associated driver genes and confer mutant progenitor cells a competitive advantage that leads to clonal expansions. The human esophageal epithelium exemplifies this phenomenon, becoming a dense mosaic of competing mutant clones by adulthood. However, the phenotypic consequence of those mutations and their possible role in carcinogenesis remains unknown. Novel bioinformatic tools for *de novo* mutant detection from single-cell transcriptomics (scRNA-seq) could potentially leverage on the wealth of publicly available data to help draw mutant cell phenotypes *in vivo*. In this study we test SComatic algorithm’s ability to identify somatic mutations in the normal, polyclonal esophageal epithelium. We analyze a public scRNA-seq dataset from a human cohort with multiple esophageal samples per donor, and an independent study in mice subjected to experimental mutagenesis where samples have been re-sequenced for validation. These unconventional experimental designs allow us to control unspecificity. We observe scRNA-seq variant calling output is heavily affected by undesired technical artifacts and germline variants, which we are able to reduce following a customized series of rational filters that enrich in somatic mutations. Final candidate mutations are then used to reconstitute clonal lineages and map them to differentiation trajectories in the UMAP embeddings. We find low read depth and sparse cellular sampling favor detection of passenger mutations and hinder driver mutant phenotypic inferences. Altogether, we showcase current limitations of scRNA-seq-derived mutation calling, while we offer methodological indications that should be considered for future studies aimed at investigating mutant clone behavior in normal polyclonal tissues from single-cell transcriptomics.

## INTRODUCTION

Mutagenesis is not a phenomenon exclusive to cancer. Recent advances in deep DNA sequencing technologies have enabled a plethora of studies reporting the accumulation of mutations in a wide variety of somatic tissues as we age (Yoshida 2026). These range from the haematopoiesis line to different epithelia of diverse renewal rates and even neurons (Jaiswal et al. 2014; Martincorena et al. 2015; Lee-Six et al. 2019; Lawson et al. 2020; Lodato et al. 2018). A paradigmatic case is the human esophageal epithelium, where in a healthy individual by middle age more than half of its surface gets colonized by mutant clones, many bearing alterations in cancer-associated genes (Martincorena et al. 2018; Yokoyama et al. 2019). Genes such as *NOTCH1* and *TP53* are found under strong positive selection. Some of these mutant clones reach a size of up to several millimeters, yet the tissue preserves an apparently normal histology. This poses the fundamental question of what the phenotypic consequences of those mutations are: how they impact progenitor cell function and whether they can contribute or not to carcinogenesis (Higa and DeGregori 2019; Acha-Sagredo et al. 2022).

To address some of these questions we and others have made use of lineage tracing in transgenic mice (Kretzschmar and Watt 2012). The mouse esophageal epithelium is a stratified squamous tissue where progenitor cells located in the basal layer regularly divide, and cells exit the cell cycle and transit through suprabasal layers as they terminally differentiate (Piedrafita et al. 2020). Based on conditional *Cre-loxP* systems, we can induce particular driver mutations in scattered single cells, associated with the expression of a fluorescent reporter, which allowed us in the past to track the dynamics of labelled mutant progenitor clones *in vivo*. We observed how some of the most prevalent driver mutations in the mouse esophagus confer a competitive growth advantage not by increasing proliferation rate but by tilting mutant progenitor cell fate towards a decreased differentiation likelihood: both *Notch1* inactivating mutants, *p53^R245W^* and *Pik3ca^H1047R^* clones exploit impaired differentiation to colonize the tissue (Alcolea et al. 2014; Abby et al. 2023; Fernandez-Antoran et al. 2019; Herms et al. 2024). Mutant clone dynamics can be more extensively studied upon mouse exposure to the mutagen diethylnitrosamine (DEN). In that context, rare scattered tumor lesions form, characterized by the expression of Krt6a, and are found frequently mutated in *Atp2a2* gene (Colom et al. 2021). However, the vast majority of the epithelium shows a normal or hyperplastic histology, despite becoming densely mutated, especially in genes involved in the Notch pathway. Mathematical modeling suggests cell differentiation imbalance reverts in most driver mutant clones due to long-term spatial competition constraints, limiting further progression into tumorigenesis (Colom et al. 2021, 2020). This phenotypic adaptation could help distinguish normal colonizing mutations from cancer promoter ones in such a complex patchwork of mutant clones.

The human esophageal epithelium differs from mouse in that it lies on stromal protrusions called papillae and cell proliferation is not limited to the deepest basal layer but extends to the first suprabasal (epibasal) layers (Grommisch et al. 2025). However, lineage tracing techniques are obviously limited for human research. A suitable method to investigate mutant cell behavior in humans is to co-capture DNA and RNA transcripts at single-cell resolution and perform paired DNA and RNA sequencing assays, a strategy that is gaining interest in the field (Olsen et al. 2025). However, it is still expensive and, in practice, experiments to date get circumscribed to a small target gene panel, except in rare occasions (Yuan et al. 2026; Prieto et al. 2025).

A tempting alternative would be to perform variant calling directly from single-cell RNA sequencing (scRNA-seq), taking advantage of the growing amount of publicly available data from this technique. Various methods have emerged in the last few years for this, yet most require support from parallel DNA sequencing for background modeling and/or variant validation (Huang et al. 2020; Liu et al. 2019). However, a recent method stands out for relying just on scRNA-seq data alone and it has been shown to outperform others: SComatic (Muyas et al. 2024). SComatic performs *de novo* variant calling at cluster-based level, exploiting background base call statistics for mutation filtering. Following pre-annotated cell populations, it is particularly robust when at least one cluster can be taken as pure non-mutated reference population, being thus well suited when comparing distinguishable tumor versus paired normal scRNA-seq clusters. However, its performance on intrinsically heterogeneous polyclonal populations such as those in adult somatic tissues has not yet been tested.

In this work we test *de novo* scRNA-seq variant calling and discuss its limitations on mutant discovery and phenotype inference in the genetically mosaic esophageal epithelium, by applying SComatic to both human and mouse data. We conveniently exploit scRNA-seq datasets from one study where multi-site biopsy sequencing was performed and another where sample biopsies were sequenced twice. These experimental designs allow us to monitor the method’s accuracy and evaluate the extent of clonal inferences on somatic evolution that can be drawn from it.

## RESULTS

### scRNA-seq calling of somatic mutations in mouse esophagus after mutagenesis

To test whether single-cell transcriptomics data is suitable to detect somatic mutations in a highly polyclonal, normal tissue, we first focused on scRNA-seq data from the esophagus of mice subjected to experimental mutagenesis (**Fig. 1A**). In this experiment, a sample was procured from the peeled esophageal mucosa of aged mice that had been exposed for 2 months to DEN along with a sample pooled from age-matched control littermates. Both samples were submitted to 10x Chromium (3’ capture) scRNA-seq alongside six other libraries corresponding to young adult control littermates which had been previously published (McGinn et al. 2021) (**Methods**). All eight libraries were re-sequenced here at a higher depth for the purpose of testing reproducibility in variant calling (**Fig. 1A**).

**Fig. 1.**
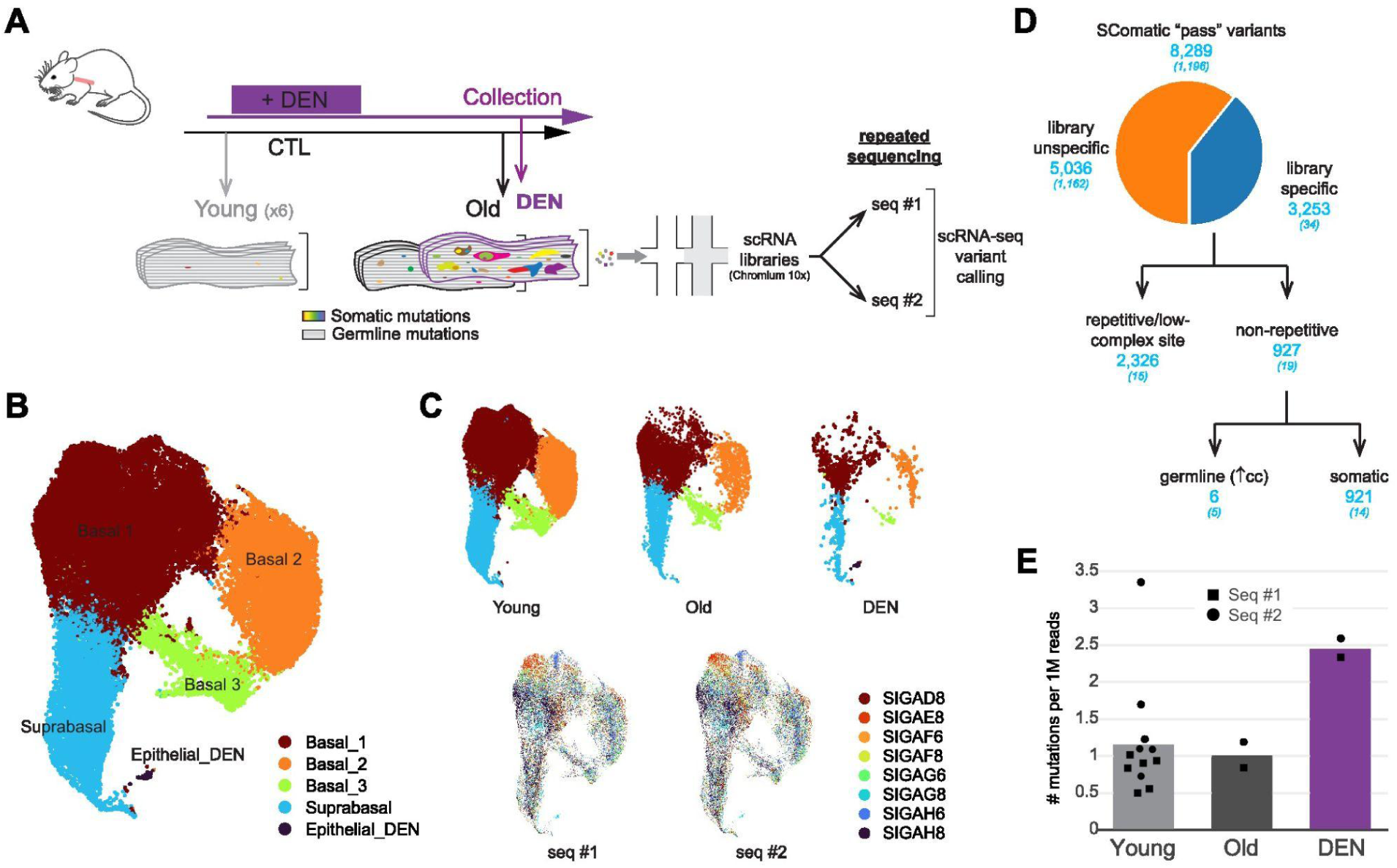
Detection of somatic mutations from mouse esophageal scRNA-seq data. (A) Experimental design describing sample acquisition and Chromium 10X libraries, which were re-sequenced by scRNA-seq. Germline variants are expected to be commonly shared across littermates and/or relatively abundant in individual libraries, unlike somatic mutations. DEN, Diethylnitrosamine. (B) UMAP-embedding of annotated epithelial cell populations in the dataset (*N* = 39,763 cells). (C) UMAP-embedding highlighting epithelial cells represented in each condition (top panel) and sequencing run (bottom panel; color coding according to library ID). (D) Customized filtering workflow used to select somatic mutations from SComatic output variants. The total number of candidate variants at each filtering step is shown (in parenthesis those that are supported by both sequencing runs). (E) Normalized mutational burden across sample conditions. The number of single-nucleotide variants (SNVs) is normalized by total read depth (1M reads) per library.

After data quality control, cellular annotation and filtering of a minor proportion of stromal cells, transcriptomes from a total of 39,763 epithelial cells were kept for analysis (**Fig. 1B**; **Suppl. Fig 1**; **Methods**). Cellular transcriptomes classified in five different clusters, four of which were evenly represented across conditions and between both sequencing runs (**Fig. 1C**). Of these, three were identified as progenitor basal keratinocytes in different cell-cycle stages and the other as differentiated, suprabasal cells based on the expression of canonical squamous epithelial markers (**Suppl. Fig 1**). Cell-cycle markers as well as trajectory inference helped to draw a neat differentiation axis along the UMAP embedding (**Suppl. Fig 1**). Intriguingly, a fifth isolated cluster was exclusive of DEN-treatment condition (reproduced in both sequencing runs). This showed an abnormal expression profile of proliferation- and differentiation-related genes, it was enriched in pathways related to stress response, immune activation, cell cycle control and cell growth programs, and it was characterized by a high expression of *Krt6a* and *Krt17*, strongly suggesting it corresponds to a subpopulation of stressed cells forming early lesions (**Suppl. Fig 1**). From here, we conclude that single-cell transcriptomes in mouse esophagus upon mutagenesis are adequate for phenotype mapping, allowing to discriminate tumor cell identities and ascribe non-tumoral individual cells to different proliferative states and fates along the differentiation trajectory.

We next applied the SComatic pipeline to detect somatic mutations in this dataset. Without prior evidence on a given cluster that could act as a germline reference population, we performed variant calling at *sample* level on each sequencing batch separately. We thus collected base count information from each library-specific BAM file and followed default SComatic parameter settings (**Suppl. Fig 2**; **Methods**). This analysis yielded a total preliminary list of 8,289 single-nucleotide variants (SNVs) between both sequencing runs (**Fig. 1D**). True somatic point mutations are generally unlikely to be recurrent across individual mice and are thus expected to be restrained to a single library, except for recurrent hotspots (Martinez-Ledesma et al. 2020). 61% of all variants were shared across multiple libraries, indicating that they either correspond to strain-related, common germline polymorphisms, sequencing errors or that they map to repetitive or low complexity genomic regions, as further evidenced by their widespread distribution across cellular barcodes (**Suppl. Fig 3**). In addition, we found 72% of the remaining 3,253 variants occurring in a single library indeed mapped to repetitive genomic regions as defined by RepeatMasker, and were thus discarded (**Methods**). In turn, a few other variants were spread through a strikingly large fraction of cells (around ∼10%) in their given sample library and were removed too as they were suspected of being mouse-specific germline or early developmental mutations (**Suppl. Fig 3**). This led to a final set of 921 candidate somatic mutations, representing just 11% of the initial list. It follows that irrespective of the sequencing run, once corrected by differences in coverage, more than twice as many mutations were found under DEN-treatment condition than in control samples. This relative increase seems modest considering the mutagenic stimulus, but it is consistent with an enrichment in somatic variants (**Fig. 1E**). Altogether, customized rational filtering is critical for *de novo* somatic variant detection from scRNA-seq, when no paired genomic data is available to build individual germline references, even though mutation discovery from scRNA-seq might be limited, as we will address below.

### Annotation and clonal features of scRNA-seq derived mutations in mouse esophagus

We followed up by evaluating the agreement between the mutations found in one and the other sequencing run (**Fig. 2A**). Surprisingly, only 14 somatic SNVs (1.5%) were supported by both datasets - whereas the highest proportion of somatic mutations (77%) were exclusively called by the second run (the one performed at a higher depth: 308 ± 104 vs. 70 ± 25 M reads / library). We speculated that this reduced overlap could be explained by the limited sensitivity due to the low average read depth that is inherent to scRNA-seq data. In fact, the average number of variant reads called per mutated cell barely exceeded the detection limit (value of 1), with just a minor but significantly higher variant depth shown by those mutations shared between sequencing runs, indicative of a relatively higher gene expression (**Fig. 2B**; **Suppl. Fig 4**). Another relevant factor is cellular sampling. For any given library, there was a high level of overlap between cellular barcodes visited by both sequencing runs, indicative of limited library size (**Fig. 2C**). We matched variants to individual cells after computing the genotype for each cell at the variant sites (**Suppl. Fig 2**; **Methods**). More than 75% of all mutations were restricted to 2-4 cells, right above SComatic detection cutoff, suggestive of sparse clonal coverage (**Fig. 2D**). Shared mutations did not show but a modest trend towards a larger (but still limited) number of cells, and no noticeable differences were seen between experimental conditions (**Suppl. Fig 4**). This is consistent with a highly polyclonal tissue, where only a small fraction of dissociated tissue cells are processed and multiple cells from a given clone are unlikely to be co-sampled for scRNA-seq (i.e. tissue cell composition is underrepresented in scRNA-seq). Therefore, the measured number of mutant cells per variant (i.e. the *apparent* clone sizes) underestimates *actual* clone sizes, and does not precisely measure the extent of mutant clonal expansions, despite providing consistent numbers between both sequencing runs for most noticeable mutations (**Suppl. Fig 4**) (Hereafter, for shared mutations, mutant cells in each sequencing run were considered as separate clones). Altogether, both transcript read depth and library size limit mutant detectability by scRNA-seq technology.

**Fig. 2.**
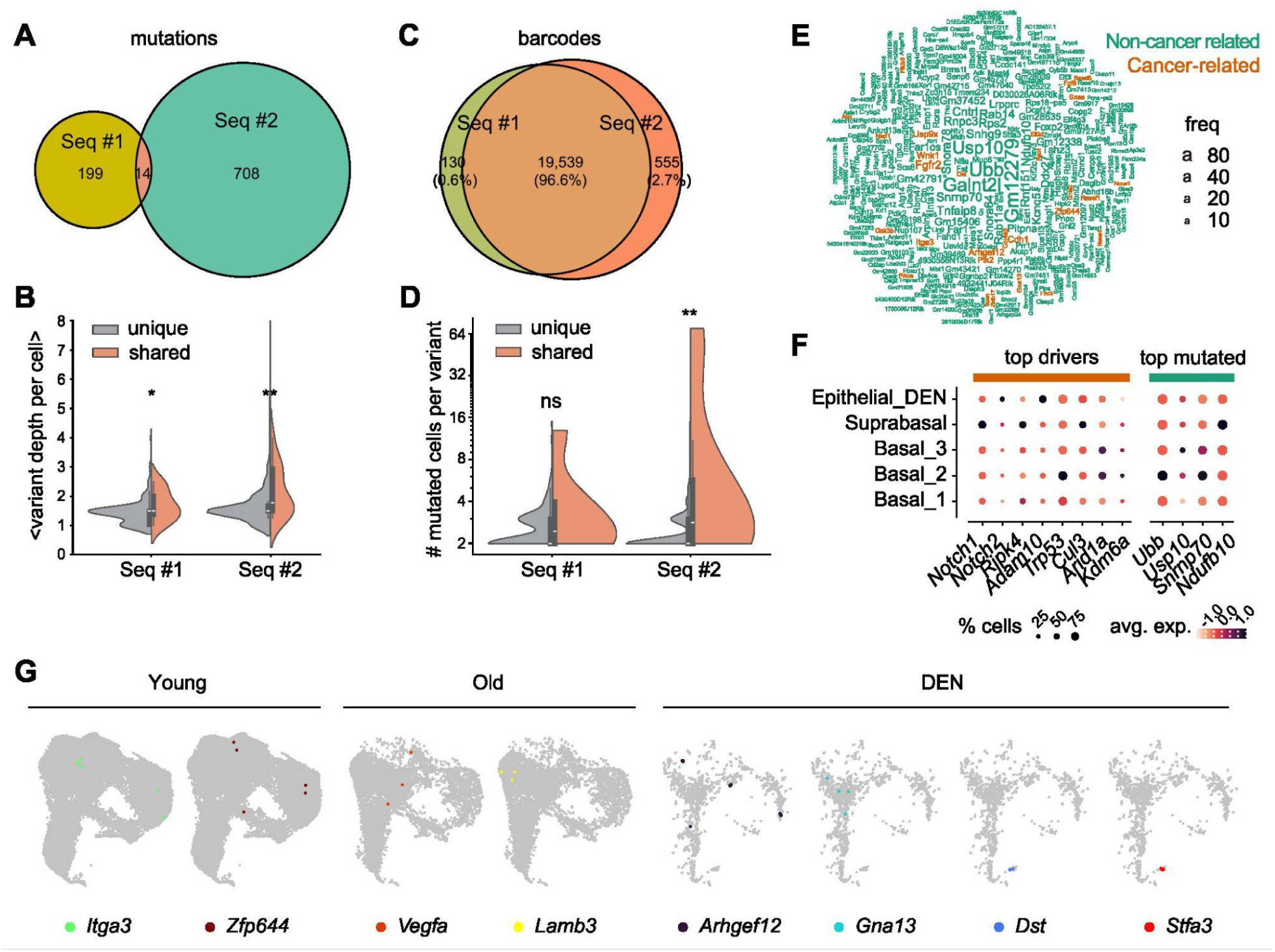
Annotation and clonal features of scRNA-seq derived mutations in mouse esophagus. (A) Venn diagram showing the intersection between the variant genomic sites found in each sequencing run (7% and 2% of variants in seq #1 and #2 are shared, respectively). (B) Distribution of the average number of variant reads (base counts) per mutated cell supporting each mutation, comparing mutations detected by a single sequencing run to those shared across both sequencing runs. Cellular statistics are divided by sequencing session (*p*-val: 0.035, 0.003 for seq #1 and #2, respectively; one-sided Mann-Whitney *U*-test). (C) Venn diagram showing the intersection between all cellular barcodes found in each sequencing run. (D) Distribution of apparent clone sizes (number of mutant cells per variant), comparing single-run called mutations to mutations shared across sequencing runs. Cellular statistics are divided by sequencing session (*p*-val: 0.100, 0.004 for seq #1 and #2, respectively; one-sided Mann-Whitney *U*-test). (E) Word-cloud of top 400 most frequently mutated genes. Names are scaled by incidence (total number of cells with variant calls for that gene). Cancer-associated genes are highlighted in orange. (F) Average normalized gene expression and detectability across epithelial single-cell transcriptomes of most prevalent driver genes found mutated in DEN-treated mouse esophagus in DNA-seq studies and top mutated genes from our analysis. Dot size indicates the % of cells where the gene was detected. (G) Examples of mutant clones found in different conditions, mapped to individual cell transcriptomes in the UMAP embeddings.

To assess biological impact, we annotated the mutations using VEP (**Suppl. Fig 2**; **Table S1**; **Methods**). Among the wide variety of mutated genes, functional analysis revealed an enrichment in genes involved in pathways in cancer (**Table S2**). However, only 83 out of the total 935 clones were mutated in cancer-associated genes (**Fig. 2E**; **Table S3**). The top prevalent variants detected in more cells occurred on high-expression, ubiquitous genes involved in housekeeping functions, such as protein ubiquitination (*Ubb* and *Usp10*), mRNA splicing (*Snrnp70*) or energy metabolism (*Ndufb10*). Intriguingly, no mutations were found in either *Notch1*, *Notch2*, *Ripk4*, *Adam10*, *Trp53*, *Cul3*, *Arid1a* or *Kdm6a*, the most prevalent driver mutant genes found in DEN-treated mouse esophagus in DNA-seq studies (Colom et al. 2020). Many of these showed modest expression values (**Fig. 2F**). Altogether, variant calling from 10x Chromium scRNA-seq might be limiting for cancer driver genes, either due to low expression levels or, alternatively, sequence biases linked to the 3’ mRNA capture technology.

Overall, mutations classified as modifier (mapped to introns or UTR regions) and (to a lesser extent) moderate impact (missense) dominated over high-impact ones (non-sense and splice-site variants), suggesting influence from short 3’-end sequence bias. In any case, impact severity showed a negligible effect on mutant clone extension or detectability, suggesting the predominance of passenger rather than driver mutations (**Suppl. Fig 5**). Yet, we found some potentially relevant mutations within the group of cancer-related genes, like a high-impact, splice-donor variant in *Arhgef12* or a 3’-UTR variant in *Gna13*, both drivers of tumorigenesis and metastasis in mouse and human cancers that were mapped to predominantly basal-cell clones under DEN condition (**Fig. 2G**). Instances of cancer-related mutants constrained to the basal proliferative compartment were observed too in CTL, exemplified by two clones in old mice with missense variants in *Vegfa* and *Lamb3*, respectively, or a pair of clones mutant for *Itga3* and *Zfp644* in young mice. Conversely, clones circumscribed to the DEN-specific tumor-like epithelial cluster were found as well, like a 2-cell clone with a high-impact, stop-gained mutation (p.R7362X) in *Dst*, a gene encoding for dystonin, where loss-of-function mutations have been reported to cause epidermolysis bullosa simplex (Groves et al. 2010), or a 5-cell clone with a missense variant (p.E45K) in *Stfa3*, encoding for cystatin A, a regulator of keratinocyte differentiation which is strongly upregulated in AP1/AP2-altered mice presenting a keratoderma-like skin phenotype (Rorke et al. 2015; Zhang et al. 2024) (**Fig. 2G**).

Despite comprising a relatively small fraction of cells, the DEN-specific cluster concentrated an increased density of mutations (*p*-val: 0.007; Fisher exact test). Moreover, among its intersecting mutant clones, the number of those that were fully embedded (with all cells annotated to this cluster) was higher than expected by chance (*p*-val < 0.0001; random permutation test), a pattern consistent with an intratumoral clonal origin (The rest showed partial overlap as they included cells from other clusters, and could presumably be interpreted as pretumoral). We explored the clonal structure across the entire dataset and found that 75% of all detected mutations fell in independent groups of cells, forming isolated clones; so did they in the DEN-specific cluster, in agreement with the high polyclonality and multiple tumor foci (**Suppl. Fig 5**). When present, colocalized mutant clones overwhelmingly defined clusters of cells with incomplete cellular overlap, attributed to the sparse read coverage per cell (**Suppl. Fig 5**). Altogether, mouse scRNA-seq-derived mutations can be mapped to some individual cell states and ascribed to broad normal or tumoral cell lineages, allowing rough phenotypic inspections, even though they are not suited for fine subclonal inferences.

### Detection of somatic mutations from human esophageal scRNA-seq data

Once we set the limitations of scRNA-seq derived mutation calling for somatic evolution studies in mice, we turned to explore mutation inference in human data. In contrast to experimental models, where a great genetic homogeneity can be assumed between littermates (most suspected germline variants in mice were indeed filtered out when restraining to single-library variants), inter-individual heterogeneity poses a challenge for discriminating germline polymorphisms in human cohorts. Hence, we selected a particular published, high-throughput dataset from the Human Cell Atlas project where separate libraries for 10x Chromium scRNA-seq were procured from 4 adjacent regions of esophagus in each of 6 adult healthy donors (Madissoon et al. 2019) (**Fig. 3A**). We reasoned that this multi-site sampling design could aid identifying and discarding individual-specific germline variants, as set below. A total of 87,947 cell transcriptomes were contained in the author’s dataset after curation (quality control re-assessment), distributed in 19 different clusters representing distinct epithelial and stromal cell populations (**Fig. 3B**; **Methods**) (Notice here processed cells corresponded to unsorted populations unlike in the mouse study). In particular, keratinocytes comprised 92% of all cells and were segregated into four distinguishable epithelial compartments: basal cells, deep suprabasal cells, (mid) stratified cells and upper-layer cells, with similar UMAP-projection patterns conserved across donors (**Suppl. Fig 6**). Cell-cycle phase inference indicated active proliferation concentrated to basal and deep suprabasal compartments, as consistent with molecular studies in human esophageal epithelium (Ferrer-Torres et al. 2022) (**Fig. 3D**). RNA velocity and pseudotime analysis confirmed the major alignment of cells along the terminal differentiation trajectory, altogether evidencing, as in mice, data suitability for phenotypic inferences on individual cell differentiation status (**Fig. 3D**).

**Fig. 3.**
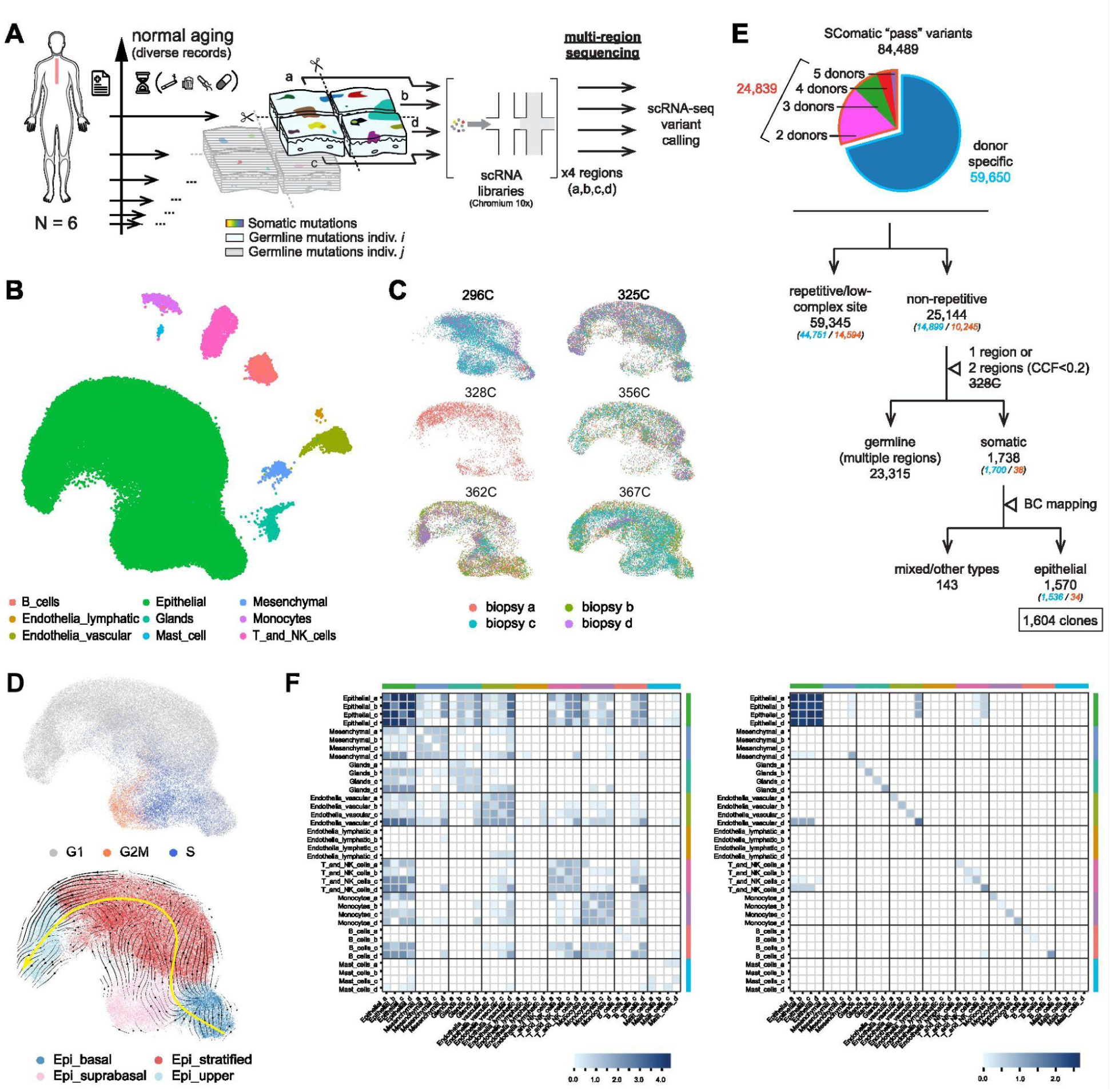
Detection of somatic mutations from human esophageal scRNA-seq data. (A) Protocol describing sample acquisition and libraries used for Chromium 10X scRNA-seq, which were procured from different adjacent regions of esophagus in each donor (4 libraries each). Germline variants are expected to be donor-specific but common across libraries procured from the different regions, unlike somatic mutations which would be localized and would encompass two adjacent segments maximum. (B) UMAP-embedding of high-quality cell transcriptomes in the dataset, highlighting the main cell-type annotations used here for variant calling (*N* = 87,947 cells). (C) UMAP-embedding subsetting epithelial cells from each donor, highlighting libraries (territories) of origin. (D) Inference on epithelial differentiation trajectory based on cell-cycle marker expression score (top panel) and RNA velocity stream vectors (bottom panel). The overlaid yellow line corresponds to Slingshot pseudotime trajectory. (E) Customized filtering workflow used to extract somatic mutations from SComatic output variants. The total number of candidate variants in each filtering step is shown (in parenthesis those repeated across various donors). (F) Heatmaps showing the frequency of co-occurrence of variants between the 36 different cell types-segments (i.e. 9 cell types x 4 segments) before (left panel) and after (right panel) filtering germline variants. Values given in log10 scale. Notice most variants encompassing 2 cell type-segments are exclusively epithelial post-filtering.

For the purpose of exploring somatic mutations, which might be present in cells encompassing various epithelial layers, we summarized the annotated cell types into main developmental lineages, so that all epithelial subpopulations were grouped together (Zhang et al. 2021) (**Fig. 3B**; **Suppl. Fig 6**). Conversely, we kept separate annotations (separate clusters) for cellular transcriptomes from the different segments (territories) of origin biopsied (**Fig. 3C**). In this manner we ran SComatic variant calling at cell-type level on each individual separately, relaxing the parameter `--max_cell_types` to 4 to account for possible variants in a given lineage potentially spreading through multiple or all biopsies from the same donor (even at the expense of retaining variants affecting multiple lineages too; base count matrices computed for every lineage-segment-specific BAM file) (**Suppl. Fig 7**; **Methods**). Overall, a significant fraction (29%) of the initial 84,489 single-nucleotide variants (SNVs) called as passed by SComatic was found repeated across various donors, raising concerns on their origin as recurrent somatic mutations (despite having applied previous SComatic filters against common polymorphisms and artefacts). No harsh, *ad hoc* filters were applied at this stage, however, given that specific downstream rational filters were devised later on (**Fig. 3E**).

In particular, 59,345 of the original variants mapped to repetitive regions or low complexity genomic sites and were excluded using RepeatMasker. From the remaining 25,144 variants, an overwhelming fraction (96.3%) were detected across multiple segments or cell types, strongly suggesting a predominance of undesired, germline and/or developmental variants (**Suppl. Fig 8**). Largest driver mutant clones in healthy human esophagus are in the mm^2^ range and are expected to be captured by a single segment or two segments maximum (when falling in between two adjacent biopsies) following manual sample trimming (Martincorena et al. 2018). Conversely, germline and early developmental variants should be retrieved in all territories except when expression levels and read depth limit detectability in one or another segment (**Fig. 3A**). Consistent with this reasoning, we observed that variants affecting multiple territories or lineages were predominantly widespread across all cellular barcodes, except for a minor fraction of those found in 2 territories which occurred at lower frequencies comparable to those present in a single segment (a fraction of mutated cells of ∼50% reflects the case of heterozygous SNPs when read depth per cell is insufficient to cover both alleles at the same time) (**Suppl. Fig 8**). For this reason, we circumscribed to variants present in one territory and those in two territories (or cell types) with a cell fraction < 20% (we removed too the variants from one donor with just one territory with high quality data). With these filters the list of candidate somatic mutations considerably decreased to 1,738 (a 2% of the original set), where the fraction of variants shared across donors became negligible (**Fig. 3E**). Importantly, variants of pure epithelial origin became predominant and potentially spurious calls involving different lineages were reduced, in agreement with an enrichment in true somatic mutations (**Fig. 3F**). All these mutations were restrained to a small proportion of cells (<10%) in their respective library (**Suppl. Fig 8**). Since clonal expansions in the esophagus localize to the highly proliferative epithelial compartment, the filtered 1,570 epithelial mutations were the ones retained for subsequent analyses. Altogether, multi-site sampling highlights the high impact of germline variants on *de novo* mutant detection from human scRNA-seq data and the need for appropriate customized filtering.

### Annotation and clonal features of scRNA-seq derived mutations in human esophagus

We analyzed the mutational characteristics across individuals (**Fig. 4A**). A variable mutation burden was observed between individual donors, with no clear correlation with age, possibly due to heterogeneous exposure histories (and/or uncontrolled technical noise). In turn, apparent mutant clone sizes remained restricted to a few cells, typically between 2-5, consistent with the limited transcript detection by scRNA-seq, as we previously reported with mice. Following VEP annotation, we found most variants were catalogued as modifier (intronic or UTR-mapped) or low impact (synonymous) (**Suppl. Fig 9**; **Table S4**). The low representation of missense and nonsense variants (of moderate and high impact, respectively) could again be biased by the 3’ capture scRNA-seq technique but the lack of a noticeable correlation between apparent clone sizes and impact level suggested the predominance of passenger mutations over driver mutations.

**Fig. 4.**
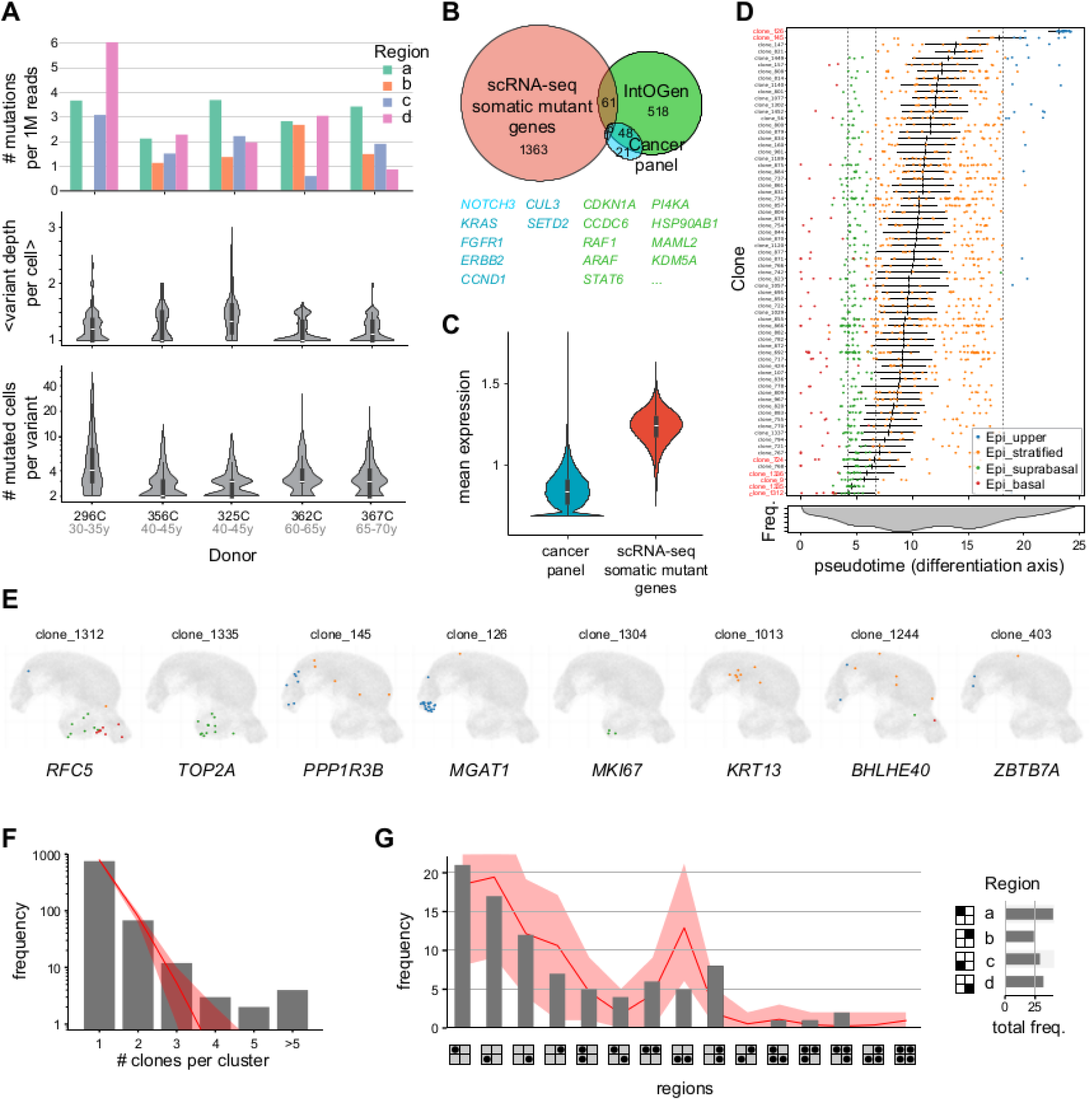
Annotation and clonal features of scRNA-seq derived mutations in human esophagus. (A) Normalized mutation burden (upper panel), distribution of the average number of variant reads (base counts) per mutated cell supporting each variant (mid panel), and distribution of apparent clone sizes (number of mutant cells per variant) (bottom panel) shown across individual donors. In the top panel, the number of single-nucleotide variants (SNVs) was normalized by total read depth (1M reads) in each library, i.e. per segment biopsied. (B) Venn diagram illustrating the intersection between the mutated genes and common cancer driver genes from IntOGen database and those in (Martincorena et al. 2018) panel. (C) Distribution of average normalized gene expression across cells for the set of genes called as mutated by scRNA-seq in comparison to cancer-associated genes from (Martincorena et al. 2018) panel (p-val < 2e-16; one-sided Mann-Whitney U-test). Only cells with non-zero gene expression values were considered. (D) Cellular composition of largest mutant clones (those detected in ≥10 cellular barcodes), displaying individual cells colored by epithelial subtype and ordered by pseudotime score (mean values with 95% confidence interval shown as errorbars). Below is the overall histogram of pseudotime scores from all 61,241 epithelial barcodes in the scRNA-seq dataset. Clones with significantly biased cellular pseudotime scores are highlighted (two-sided Mann-Whitney *U*-test, with B-H correction for multiple testing). (E) Examples of mutant clones found in the human cohort, mapped to individual epithelial cell transcriptomes in the UMAP embeddings. Mutant cells are color-coded according to epithelial subtype, as in (D). Affected genes are shown underneath. (F) Frequency of individual mutation co-localization across cells, showing involvement in increasing size clusters. Cluster sizes are given by the number of individual mutations sharing at least one overlapping mutant cell in common. Best-fit Poisson distribution (expected rate: λ=0.201) is shown overlaid, with 95% uncertainty margins given by the actual sampling error (bootstrapping) (**Methods**). (G) Distribution of the 89 mutant clusters grouping two or more clones across spatial territories, detailing frequencies of clusters spanning one, two or multiple territories (Overlaid is the random expectation obtained by permutation and linkage of coherent clones given individual clone location frequencies, with 95% uncertainty margins). The overall cluster frequencies on each of the four segments biopsied are shown in the legend.

The intersection with existing curated human variant databases allowed us to validate the nature of some inferred mutations (**Suppl. Fig 9**; **Methods**). Around 30% of the mutations were both reported in dbSNP database and found with at least a minimal allele frequency (≥ 1e10^-6^) in human populations (1,000 Genomes or gnomAD projects) being thus suspicious of unfiltered germline variants. The rest comprised a range of mutations (∼10%) with strong evidence for being somatic (i.e. annotated in COSMIC and/or dbSNP databases and with no reported incidence in healthy human cohorts) and a large fraction (∼60%) of unknown variants, a common feature in ultradeep or single-cell DNA sequencing studies (Lawson et al. 2025). Unknown variants shared similar apparent clone sizes and gene structure mapping characteristics as known somatic mutations, unlike suspicious germline variants, arguing on their somatic origin (**Suppl. Fig 9**). Altogether, around ∼70% of filtered variants could be taken as trustable somatic mutations, and hence used for subsequent analyses.

We next explored gene annotations among mutations predicted as somatic. Pathway analysis revealed an enrichment in housekeeping functions such as protein metabolism and cell adhesion (**Table S5**). Mutations in widespread-expression genes such as *TMTC3* and *PPP2CA* (encoding post-translational modification enzymes), *EEF1A1*, *RPL38* and *EIF2S3* (encoding translation machinery components) or *ACTB*, *WASF2* and *NDRG1* (involved in actin cytoskeleton reorganization and/or cellular trafficking) were among the ones detected in more cells (**Suppl. Fig 10**). In contrast, very few mutations mapped to cancer-associated genes. For instance, only seven genes from (Martincorena et al. 2018) panel were found mutated: *NOTCH3*, *KRAS*, *FGFR1*, *ERBB2*, *CCND1*, *CUL3* and *SETD2* (all of them in intronic or 3’ UTR regions) (**Fig. 4B**). It follows that cancer-associated genes showed very low expression values compared to other genes in the scRNA-seq dataset (**Fig. 4C**; **Suppl. Fig 10**), allegedly limiting detection of potentially driver, pathogenic mutations from single-cell transcriptomics.

Following barcode mapping, we next explored mutant clone phenotypes. Largest detected clones commonly encompassed cells from all epithelial compartments, assigned to a broad spectrum of differentiation states (pseudotime scores), consistent with passenger mutation status and an unaltered differentiation program (**Fig. 4D**). Some clones, however, were significantly enriched in one or another epithelial subtype (**Fig. 4E**). We found modifier-class mutations mostly restrained to basal or proliferative keratinocytes for genes such as *RFC5*, *TOP2A* and *ITGB3BP*. Conversely, mutations affecting *MGAT1*, *PPP1R3B* or *FDFT1* gene transcripts were seen predominantly on terminal differentiated cells. Indeed, the former are common markers of keratinocyte proliferation, whereas the latter play a role in epithelial barrier function and are thus upregulated during terminal differentiation. From here, we interpreted that clones with significantly biased cellular pseudotime scores did not reflect driver mutations altering the normal differentiation program as much as the case of variants on genes with a subtype-specific expression pattern and different transcript detectability across epithelial subpopulations. On the other hand, we looked for mutations imprinting potential specific gene expression changes, either on their own transcript (for modifier-class variants) or downstream effectors (for those very few high- or moderate-impact mutations on transcription factors): we found little evidence due to the sparse scRNA-seq data and limited clone sizes (**Suppl. Fig 10**) (Jia and Zhao 2019). Altogether, human esophageal scRNA-seq derived mutations were dominated by rare, passenger variants on detectable transcripts, heavily conditioned by gene expression levels.

Finally, we analyzed the subclonal structure across donor biopsies. Most mutations developed as independent clones, with just 11% of cases consisting of small clusters of cells sharing more than one mutation (**Fig. 4F**). Individual mutant clones were evenly distributed across spatial territories, and albeit showing incomplete barcode overlap, when present, clusters remained predominantly restricted to cells from one or maximum two biopsies, as expected according to localized events given individual clone abundances (**Fig. 4G**; **Suppl. Fig 10**). In summary, despite the limited sampling and modest detection power, scRNA-seq derived clonal structure was consistent with a scenario of widespread, random isolated mutational events.

## DISCUSSION

*De novo* mutation calling from scRNA-seq entails both technical and methodological challenges. SComatic emerged as a reliable and robust tool for this purpose. Computing variant statistics at cell-type (cluster) level and incorporating in-built beta-binomial validation testing and various filters, it was reported to outperform other popular variant calling methods (Muyas et al. 2024). However, regardless of these improvements, the specificity and sensitivity in the detection of somatic mutations has yet remained a matter of debate (Wiens et al. 2024; Marot-Lassauzaie et al. 2024). In the original publication, around 40-70% of scRNA-seq derived variants were also detected by whole exome sequencing (WES). Conversely, this figure only represented 2.5% of the total mutations called by WES (∼a half when restricting the scope to sites with sufficient coverage in both WES and scRNA-seq). Indeed, with a variant calling cutoff for WES of x10 site coverage on matched normal samples, 1/1,000 of all rare heterozygous SNPs would be expected to be overlooked, so inflation due to the contribution of unfiltered germline variants cannot be ruled out. Here, we control against germline polymorphisms in scRNA-seq mutation inference by comparing SComatic output variants between mouse littermates or across several biopsies of each individual donor in a human cohort. It follows that the vast majority of primarily acceptable variant calls in scRNA-seq data indeed correspond to unfiltered SNPs or amplification/sequencing artifacts common across libraries. A second source of false positives comes from variants mapped to repetitive or low-complexity genome sites. SComatic developers recommend intersecting output candidate mutations with RepeatMasker and indeed we observe this should be a requisite. It follows that in the esophagus data just 11% of the originally proposed variants in mice and 2% in humans can really be taken into consideration as candidate mutations. These aspects justify the importance of devising customized filtering steps on scRNA-seq variant calling.

Can we trust the final set of variants that we obtain upon filtering as somatic? Resorting to annotations from databases of known human variants, we find that 1/3 of the final set might still correspond with unfiltered SNPs in our analysis on human. This figure would be more difficult to estimate in mouse, but it is striking that a significant number of the inferred mutations were found in the normal esophagi from control adult mice. Healthy experimental mouse esophagus has been genotyped before at high depth (Colom et al. 2020), but certainly not at single-cell resolution to consider the possible contribution of passenger mutations restrained to a few cells. Nevertheless, it is appealing that the apparent mutation burden in control mice constitutes around 1/3 of that found in mice that received mutagen. One can speculate that perhaps the majority of those variants under control conditions are in fact unfiltered germline variants or technical artifacts and just a fraction of those variants in DEN mice can be trusted as somatic. Inspecting non-epithelial cells could be a possible strategy to further filter against spurious variants on ubiquitous genes when dealing with scRNA-seq data from unsorted populations. Regardless of these considerations, several elements can be taken in support of an improved somatic mutation content among the filtered variants. First, despite the sample size limitations, a higher mutation burden is seen in mouse esophagus upon mutagenesis than in age-matched controls (indeed, presumably this ratio would be comparatively higher and more realistic if a number of unfiltered germline variants were to be substracted). Second, in humans, final proposed variants appear minimally shared between individuals, unlike discarded suspicious germline variants or artifacts found across territories. Third, upon filtration, candidate human variants affecting cell populations from two different territories or biopsies get mostly circumscribed to cells from the same lineage (in particular affecting the epithelial population, which shows a high turnover in the esophagus). Finally, a consistently low variant allele cell fraction (VACF) is reported across candidate mutations annotated as unknown and those present in COSMIC, lower than that for pre-annotated human SNPs. We are aware that none of these features rules out an additional contribution from spurious variants. However, all of them point to an improved specificity in somatic mutant detection upon our customized filters. One could argue on the benefits of contrasting scRNA-seq variants with paired DNA-seq data, but standard shallow whole genome sequencing (WGS) or WES would be of little use for mutant detection in a highly polyclonal tissue (Menon and Brash 2023; Adhikari et al. 2025; Lawson et al. 2025). Validating the candidate somatic SNVs found in scRNA-seq by ultradeep DNA-seq or scDNA-seq is out of the scope of this survey.

The apparent absence of driver mutations in this study can be due to different reasons. First, scRNA-seq variant calling suffers from low sensitivity. Our experimental design allowed us to estimate a 2-7% overlap in variant recovery rate between independent sequencing runs of the same libraries in mouse (despite high concordance in the cell barcodes recovered). Cancer-related genes exhibit heterogeneous expression levels and we show scRNA-seq mutation discovery is unavoidably biased towards variants on constitutive, high-expression genes, which are most easily detected. These would be subject to purifying selection insofar as they exert housekeeping functions (Zhang and Li 2004). Even within this category, scRNA-seq data is sparse in terms of read counts per cell at individual gene level. Overall, we detect ∼1 mutation every 3 single-cell transcriptomes in our analysis in human esophagus, when adult individuals accumulate hundreds to thousand mutations per cell in genomic studies in this tissue (Martincorena et al. 2018). Yet, if we circumscribe to those genomic regions with sufficient read coverage we obtain an estimated mutational burden of 0.3-1 mutations per Mb, which is in agreement with somatic evolution evidence (Bridge et al. 2025), further supporting the low false positive rate and expected somatic origin of the filtered variants. Adopting strategies that preamplify specific transcripts of interest, such as the ones used for clonal barcode tracking, could be instrumental for enhancing driver mutant detection (Yang et al. 2022; Weber et al. 2025).

Secondly, we chose scRNA-seq data coming from 10X Genomics Chromium Single Cell 3’ platform, for being the most widely used in the field. This technology captures a subset of the transcriptome, specifically focussing on the 3’ ends of mRNA (poly-A tail capture), rather than providing full (and homogenous) coverage of the entire transcriptome. It follows that a high fraction of variants that we find are annotated to UTR regions. More surprising is the relative abundance of variants affecting intronic or even intergenic regions. However, this is a common feature with datasets in the original SComatic publication. The existence of a high proportion of variants mapping to intergenic regions was further supported by a pan-tissue analysis on scATAC-seq (Muyas et al. 2024), arguing against scRNA-seq technology-specific artifacts. Off-target, intergenic reads are common in 10x 3’ data and frequently originate from transcribed functional elements in non-coding regions (He et al. 2024). Circumscribing to those mutations falling on coding regions, we find more missense than synonymous variants (a pattern that consistently differs from filtered SNPs) and there are very few nonsense variants, consistent with the fact that truncated transcripts are often prematurely degraded via nonsense-mediated decay.

Wrapping up around the point of driver mutation discovery on cancer-related genes, on one hand, loss of function mutations in tumor suppressors would be difficult to be detected by transcriptomics as they would often result in a silent or very low signal. On the other hand, detection of activating (missense) mutations in oncogenes by scRNA-seq might be favored when these lead to over-expression. However, the mRNA 3’ end biased capture method would limit their detection in target genes with long 3’ UTRs or when driver mutations fall far upstream an oncogene’s transcript, e.g. this is the case for main driver mutations on *TERT* or *KRAS* (Gasper et al. 2022; Kaya et al. 2021). It would be interesting to test scRNA-seq variant calling with alternative, full-transcript technologies, such as Smart-seq2 (Dondi et al. 2025).

Mapping mutations to cellular barcodes allows exploration of tissue subclonal structure. The frequency of events of co-localization between mutant clones approximately conforms to a Poisson distribution both in mouse and human, which suggests a landscape of randomly scattered clones. According to this model, the probability of observing two or more independent events co-occurring in space is as small as 1.6e-5, which would correspond to a total sampling area of ∼200,000 cells given an average clone size of 3-4 cells. This extension is consistent with a mm-size biopsy and it is one order of magnitude higher than observed scRNA-seq library sizes, supporting the argument of sparse clonal representation after sample dissociation and library preparation constraints. Not surprisingly, co-localizing mutations group into vague clusters of incomplete cellular overlap that preclude subclonal lineage inferences. Nonetheless, despite this coarse level of granularity, captured mutant clones in scRNA-seq still look overwhelmingly small compared with the extension of drivers found by DNA-seq studies (Martincorena et al. 2018; Yokoyama et al. 2019). On one hand, this result might feel discouraging but can be expected: our data suggests that the vast majority of hits correspond with passenger mutations, which also constitute the bulk of variants in single-cell or ultradeep DNA-seq approaches (Kumar et al. 2020; Lawson et al. 2025). On the other hand, to date, there have already been successful examples where high frequency scRNA-seq variants are used at cell population level for sample genotyping, donor ancestry inference, identification of admixed samples and broad cell lineage tracing (Heaton et al. 2020; Ramazzotti et al. 2022; Dou et al. 2024). We believe improved confidence in passenger detection provides encouraging opportunities for detailed lineage reconstitution in human tissues with the advent of deeper and/or higher spatial resolution transcriptomic approaches (Prieto et al. 2025).

In conclusion, variant calling from short-read scRNA-seq data is to date an immature endeavor that faces both specificity and sensitivity issues. By leveraging on unconventional experimental designs involving multi-region sampling and repeated library sequencing we reveal fundamental limitations to mutation discovery from scRNA-seq data in an adult somatic mosaic tissue, showcasing a set of filters that can be applied to alleviate unspecificity issues. Although these procedures might not be readily translatable to standard RNA-seq datasets, they offer a whisper of caution on biological inference from scRNA-seq variant calling. At the same time, elaborations on similar cell type- and region-based filters could be devised to validate somatic mutation inference and test clonal structure in spatial transcriptomics data. As deeper transcriptomics approaches become affordable and spatial technologies mature, we believe our methodology including rational variant filtering, clonal structure reconstitution and visualizations can serve as a benchmark for future studies aimed at mapping clonal mutations to phenotypes.

## METHODS

### Data access

Mouse single-cell transcriptome data originated from (EpCam+/CD45−/DAPI−) sorted esophageal cells described in (McGinn et al. 2021), where pooled libraries were generated and processed using 10x Genomics Chromium technology (Single Cell 3′ v3 chemistry) (see experimental procedure therein). Here, scRNA-seq raw FASTQ files were downloaded from the public ArrayExpress repositories for the following conditions: (i) Young adult control mice of P70 days of age from the original publication (i.e. 10w old; here named “Young” condition), comprising 6 different pooled samples originated from N=3 mice each (libraries SIGAF6, SIGAG6, SIGAH6, SIGAD8, SIGAE8 and SIGAF8). These data were downloaded from the public ArrayExpress repository of the original publication, with accession code E-MTAB-8662 (https://www.ebi.ac.uk/biostudies/arrayexpress/studies/E-MTAB-8662). (ii) Pooled samples from 50w old adult control mice (“Old” condition) and (iii) age-matched diethylnitrosamine-treated mice (“DEN” condition) (libraries SIGAH8 and SIGAG8, respectively), originally processed along with those samples in (McGinn et al. 2021) and with raw FASTQ data disclosed in public ArrayExpress repository E-MTAB-17231. All libraries were rerun in two sequencing sessions at different coverage (70 ± 25 vs. 308 ± 104 M reads / library) on Illumina NovaSeq SP and S2 type flow cells: SLX-17937 and SLX-18123, respectively, hereafter referred to as seq #1 and seq #2. Data from all libraries were analyzed separately for seq #1 and seq #2, unless stated otherwise. DEN mice received diethylnitrosamine (DEN, Sigma-Aldrich; N0756) at 40 mg/L in sweetened drinking water for 24-hours, 3 days a week (Monday, Wednesday and Friday) for eight weeks, before returning to normal sweetened water upon DEN cessation until collection at 6 months post-treatment (Skrupskelyte et al. 2026).

Regarding human scRNA-seq, data from (Madissoon et al. 2019) was downloaded from HCA Data Portal (https://explore.data.humancellatlas.org/projects/c4077b3c-5c98-4d26-a614-246d12c2e5d7) following download instructions. Only files referring to esophagus were selected: FASTQ sequencing files, BAM alignment files, and one RDS file with the author’s processed Seurat object. Data consisted of 6 individuals with 4 libraries per donor acquired from 4 adjacent regions of esophagus that were dissociated and processed for 10x Genomics Chromium scRNA-seq (Single Cell 3′ v2 chemistry) following different cryopreservation times: 0h, 12h, 24h or 72h. This period did not affect single-cell transcriptome quality according to the original publication, so these segments were here referred to as region a, b, c and d, respectively, for simplicity. Data from segment b in Donor 296C was missing, making a total of 23 libraries. File metadata and supplementary tables from the HCA paper (Madissoon et al. 2019) were thoroughly inspected and processed to correctly map raw scRNA-seq files (FASTQs) to uncoded donor info and alignment BAM files for downstream analyses (**Table S6**).

### Sequence quality control and alignment

Quality control was performed over FASTQ files using FastQC v0.12.1 and a summary report for all files was generated with MultiQC v1.21. All mouse and human sequencing data passed the quality control check and were kept for subsequent analysis, except data from three biopsies (segments b, c and d) in human donor 328C, which were discarded due to low base quality (<20 Phred score extended for most of their length).

Short-read aligner STAR version 2.7.11a was used on both datasets (https://github.com/alexdobin/STAR) (Dobin et al. 2013). STARsolo options were chosen to enable downstream RNA velocity analyses (“Velocyto” parameter along with “Gene” and “GeneFull” parameters). The resulting RNA count matrices thus contained spliced and unspliced counts. The reference genomes used were GRCm38.p6 from Ensembl version 102 for mouse; and GRCh38.p14 from Ensembl version 110 for human. Both were downloaded via FTP, and previously indexed with STAR. The obtained BAM files after calling STARsolo were indexed with samtools.

### scRNA-seq data analysis

RNA count matrices from mice were preprocessed using barcodeRanks from R package DropletUtils to filter out empty droplets generated during scRNA-seq. Briefly, “gene”, “spliced” and “unspliced” matrix counts were collected to create a knee plot of total UMI counts per barcode in ranked order. A fixed cutoff of 1,000 spliced counts was chosen based on visual inspection. Filtered count matrices were imported and merged into a single Seurat object for downstream analyses.

Normalization and scaling were applied by SCTransformation on spliced assay, followed by PCA dimensional reduction keeping 17 components, tSNE reduction, and UMAP embedding (Becht et al. 2018). k-Nearest Neighbors and Leiden algorithm (Traag et al. 2019) were used for clustering, testing different resolutions ranging from 0.1 to 1.2 in steps of 0.2. A final resolution of 0.2 was selected for representing the lowest one that preserved stability and consistency, as supported by clustree R package. Cell-type annotation was carried out based on differential gene expression analysis, cell cycle phase inference and established cell-type specific markers, with reference to protein databases and EMBL-EBI Cell Expression Atlas. Clusters 7 and 4, representing fibroblast cells and an heterogeneous population of epithelial cells with a high mitochondrial gene content (>25%), respectively, were removed from the original dataset to obtain a curated epithelial subset object. This was re-normalized and re-clustered, following similar procedure as above (res. 0.2), yielding a final set of 5 subepithelial clusters that were ultimately annotated. A coarse-grained level of annotation where all epithelial cells were exclusively labelled according to LibraryID and SequencingID info was used for SComatic variant calling (see below).

For human analyses, the processed Seurat object from (Madissoon et al. 2019) was loaded from author’s available RDS file, just removing barcodes from those 3 libraries with unmet sequence quality (see above), keeping author’s predefined clusters, annotations and UMAP embedding as in their latest version from July 2020 (https://www.tissuestabilitycellatlas.org/oesophagus), except when otherwise indicated. A coarse-grained annotation of 9 major cell types combined with RegionID was chosen for SComatic variant calling, whereas a subset restricted to the four epithelial clusters (labelled Epi_basal, Epi_subrabasal, Epi_stratified and Epi_upper) was procured for trajectory inference (see below).

### Trajectory inference

For RNA velocity, STARsolo spliced and unspliced matrices were processed on filtered cells using Seurat 4.3.0 and SeuratObject 4.1.3. In the case of the human data, our calculated spliced and unspliced count matrices were matched and merged as new assays (slots) into author’s epithelial Seurat object (subset as explained above). Only counts from Ensembl ID-matching features and matching epithelial barcode IDs were incorporated so as to restrain to filtered annotated cells that had been published (Madissoon et al. 2019). After Seurat processing and epithelial cell subsetting, mouse and human Seurat objects were converted to scanpy’s H5AD file format with the R package SeuratDisk 0.0.0.9021. RNA velocity estimation was then performed on epithelial cells using the “dynamical” model from scVelo 0.3.1 with default parameters (Bergen et al. 2020), with resulting stream vectors being projected onto the UMAP embedding with Python scanpy v.1.9.6.

Linear pseudotime trajectory inference was performed on epithelial cells with R package Slingshot 2.2.0 with default parameters (Street et al. 2018) once the filtered Seurat object was converted into a SingleCellExperiment object. Individual cell-assigned pseudotime scores were used to estimate cell differentiation status.

### Somatic variant calling from scRNA-seq

Somatic variant detection from scRNA-seq data was performed at cell type level using SComatic package (Muyas et al. 2024), following specifications from the GitHub repository (https://github.com/cortes-ciriano-lab/SComatic) (**Suppl. Fig 2 & 7**). Briefly, for mice, SComatic was run independently on data from each sequencing session (seq #1 and seq #2), taking scRNA-seq library-specific alignment (BAM) files (corresponding to Seurat’s LibraryID cellular annotation level) as elements for base counting (SComatic steps 1 & 2). The recommended parameters were used except for the minimum base quality permitted, which was set `min_bq = 30` to be more stringent on base quality. For downstream statistical testing and filtering (SComatic steps 3 & 4), the minimum number of cells supporting the alternative allele to consider a mutation was set default (`min_ac_cells=2`), leaving `max_cell_types = 1` to ensure mutations were only considered if confined to a single sample. This parameter was later relaxed to assess potential mutations affecting various libraries (Notice library was here considered as cell type for SComatic purposes; at this point, we observed that splitting data into epithelial subtypes unnecessarily complicated statistical analysis). For human, SComatic was run independently on data from each donor. Library-specific BAM files, each corresponding to a single donor biopsy, were further splitted into cell-type specific BAMs, informed by the shallow annotation level on cellular barcodes set on the Seurat object (see above), so that base count info was procured for each of the 9x4 major cell type-RegionID combinations before statistical calling. Again, recommended parameters were used, except `min_bq = 30`. We kept `min_ac_cells=2` as default, and set `max_cell_types = 4`, to enable detection of mutations affecting a given lineage but potentially extending to up to the 4 adjacent biopsies (at the expense of capturing artifacts affecting different lineages, e.g. distinct populations annotated as Epithelial_a and Mesenchymal_a) (Notice the MajorCellType_RegionID annotation was here taken as cell type for SComatic purposes). Downstream optional filters were applied to both datasets to remove RNA editing sites and common germline polymorphisms and recurrent artefacts through the intersection with a panel of normals (PoN), following SComatic guidelines. SComatic variant calling outcome was obtained in the form of 6 donor-specific .tsv files (2 .tsv files, each specific of a sequencing session, in the case of mice).

Repetitive regions were filtered out as recommended by SComatic authors, using the BED file provided by SComatic for human data. As this option was not readily available for mouse genome, in that case we downloaded the annotated track of repetitive elements from the official RepeatMasker website (https://www.repeatmasker.org/) (v.4.0.6). Based on the documentation, this annotation includes interspersed repeats and low-complexity DNA sequences such as SINEs, LINEs, and LTRs. We extracted the genomic coordinates of these regions, removed header lines, and converted the data to a BED format. *`bedtools intersect -v`* option was used to filter the SComatic output files excluding variant calls falling within repetitive elements. Final variants from SComatic .tsv output annotated as PASS were subsequently filtered as detailed in Results, considering their associated annotated features (SComatic *CCF* parameter, referred to the fraction of mutated cells supporting a variant (*Cc*) relative to the total number of cells with detectable on-site read coverage (*Nc*) was here renamed as variant allele cell fraction (VACF); SComatic *DP* parameter stands for the variant read depth).

### Estimation of mutational burden

The mutational burden in the epithelial population was estimated by dividing the total number of filtered variants by the number of genomic sites with sufficient sequencing depth in the scRNA-seq data. For this, `samtools depth` was applied on epithelial cell-type specific BAM files taking a minimum base quality of 20 (1% base error) and base positions with at least one read were summed up for epithelial cells in the given library.

### Variant annotation

Filtered variants from the final mutation .tsv files were mapped to scRNA-seq barcodes by running SComatic’s SingleCellGenotype.py script (which re-computes the genotype for each cell at the variant sites). These files were then converted to standard .vcf format by means of a customized script that allocated SComatic annotations into an ‘INFO’ column, organized as a chain of semi-colon separated tagged values. Among others, the specific cellular barcodes where a given variant was found were provided, along with donor and cell type-segment info (or library and sequencing session ID in the case of mice). Ultimately, Clone IDs were included in the ‘INFO’ tab field to commonly identify cellular barcodes with variant reads mapping to the same site in the same donor (or in the same library and sequencing session in the case of mouse). Ultimately, a merged final .vcf file was produced including all relevant information from each complete dataset.

Functional variant annotation was performed on .vcf files with Ensembl’s Variant Effect Predictor (VEP), with --everything option, selecting the appropriate VEP version for each of the assemblies (102 for mouse and 111 for human).

Only annotations on canonical transcripts were considered for analysis, except when stated otherwise. Genes matching KEGG_PATHWAYS_IN_CANCER (mmu05200 entry) and/or the panel of 192 frequently mutated genes from (Colom et al. 2020) were considered as cancer-associated in mouse. In turn, genes matching KEGG_PATHWAYS_IN_CANCER.v2024.1.Hs (hsa05200 entry) and/or the panel of 76 frequently mutated genes from (Martincorena et al. 2018) were considered as cancer-associated in human.

Mutations in human were catalogued as germline, somatic or unknown according to existing evidence provided in the VEP “Existing_variation” field. dbSNP entries (rs#) with a reported allele frequency ≥ 1E-6 in any of the human populations (“AF” or “MAX_AF” fields) were considered germline, whereas other variants with pre-existing evidence were considered somatic.

### Somatic variant association into mutant clusters

Mutant clusters were defined as groups of variants sharing at least one common barcode, and hence as a measure of clonal co-localization. Due to limited read depth, most frequently, clustered clones showed incomplete cellular overlap. Clusters were given specific IDs and cluster structures were illustrated with nVennR package. The number of variants attributed to a given cluster was referred to as ‘cluster size’. Individual clones seen in isolation (with no shared barcodes) were considered as independent 1-size clusters for analysis.

### Functional enrichment analysis

Pathway enrichment analysis was performed using the g:GOSt tool from g:Profiler web server (https://biit.cs.ut.ee/gprofiler), taking KEGG, Reactome and WikiPathways databases as ontologies. Only annotated genes were considered as the background set. Top significant terms were selected, based on *p*-value, taking 0.05 as significance threshold following false discovery rate (FDR) correction by g:SCS algorithm method.

### Statistical analysis

Random permutation tests were customized for different scenarios: (i) Simulated clones of sizes equal to those detected in Epithelial_DEN cluster in mouse were built by random sampling (without replacement) cellular barcodes from the DEN library, and used to estimate the probability that a number *n* of clones fall fully embedded within the Epithelial_DEN cluster (bootstrapping with 1M repeats). (ii) A random permutation test was performed on apparent clone sizes and functional impact annotations (VEP) to assess whether high-impact variants show higher clone sizes than expected by chance. For this, functional impact labels were randomly permuted across clones 50,000 (mouse) or 100,000 times (human), followed by contingency testing. (iii) Synthetic mutant clusters of sizes equal to those seen in the human dataset were simulated in decreasing size order, randomly drawing and grouping (without replacement) individual clones with territory overlap (only consistent clones sharing at least one region in common were considered groupable): frequencies of cluster-associated aggregated regions were computed from 1,000 repeats (bootstrapping).

A least-squares fit to a Poisson distribution (with parameter λ) was performed on cluster size frequencies to simulate the probability of co-occurrence of independent clonal events. Since no data was available for the *k* = 0 index, Poisson *P*-values were renormalized to *1-P_cum_* from *k* = 1 onwards for fitting purposes. Best-fit Poisson distribution was provided with 95% uncertainty margins, reflecting the actual sampling error given the total number of clusters detected, *m*. These margins were estimated by repeated (1M repeats; bootstrapping) random resampling of *m* clusters from a multinomial with the best-fit Poisson expected *P*-values.

Group comparisons were performed using non-parametric tests: Kruskal–Wallis test and Mann–Whitney *U*-test, with significance threshold α = 0.05. Fisher exact test and Pearson Chi-spared test (for large samples) were used for contingency tests on count frequencies across categories. Where appropriate, Benjamini-Hochberg (B-H) correction was applied for multiple testing, considering significance on adjusted *p*-values.

## Data availability

scRNA-seq data from this study has been deposited in ArrayExpress under accession code E-MTAB-17231. Processed Seurat objects will become available upon publication.

## Code availability

Code generated for scRNA-seq data analysis, variant calling and somatic mutation inference from this study are available in Github: https://github.com/gp10/scRNAseq_mutClones.

## Acknowledgements

We thank Isidro Cortes-Ciriano and Francesc Muyas for their technical support and discussion on SComatic usage. We also thank Isidro Cortes-Ciriano, Francisco X Real, Jaime Martínez-Villarreal, Phil H Jones for their critical comments on the manuscript. This work was supported by Grant PID2023-148444NB-I00 funded by MICIU/AEI/10.13039/501100011033 and by ERDF/EU. In addition, this work was supported by grants to M.P.A. from the Wellcome and The Royal Society (105942/Z/14/Z and 105942/Z/14/A). This research was funded in whole, or in part, by Wellcome (203151/Z/16/Z, 203151/A/16/Z) and the UKRI Medical Research Council (MC_PC_17230), and core support grant for Cambridge Stem Cell Institute Discovery Research Wellcome Platform Discovery Research Platform for Tissue Scale Biology (226795/Z/22/Z). A.M.A. acknowledges support from a FPU24 fellowship from Spanish Ministry of Science, Innovation and Universities and a Research Assistant fellowship from Community of Madrid, cofunded by FSE+/EU. S.V.N. is supported by a FPU23 fellowship from Spanish Ministry of Science, Innovation and Universities. G.S. was funded by the Medical Research Council (MR/P019013/1) and Worldwide Cancer Research (23-0063) to M.P.A. H.A. was supported by the Leverhulme Trust (RPG-2023-136) to M.P.A.

## Author contributions

AMA, DGM and GP performed scRNA-seq data processing, variant calling and computational analyses. SVN performed functional enrichment. MRR helped in scRNA-seq analysis and statistical analysis. GS, HA and MPA contributed with experimental scRNA-seq data. GP supervised the work. AMA, DGM and GP designed the work and wrote the first draft. All authors contributed to discussion and in the elaboration of the final manuscript.

## SUPPLEMENTARY FIGURES

**Suppl. Fig. 1.**
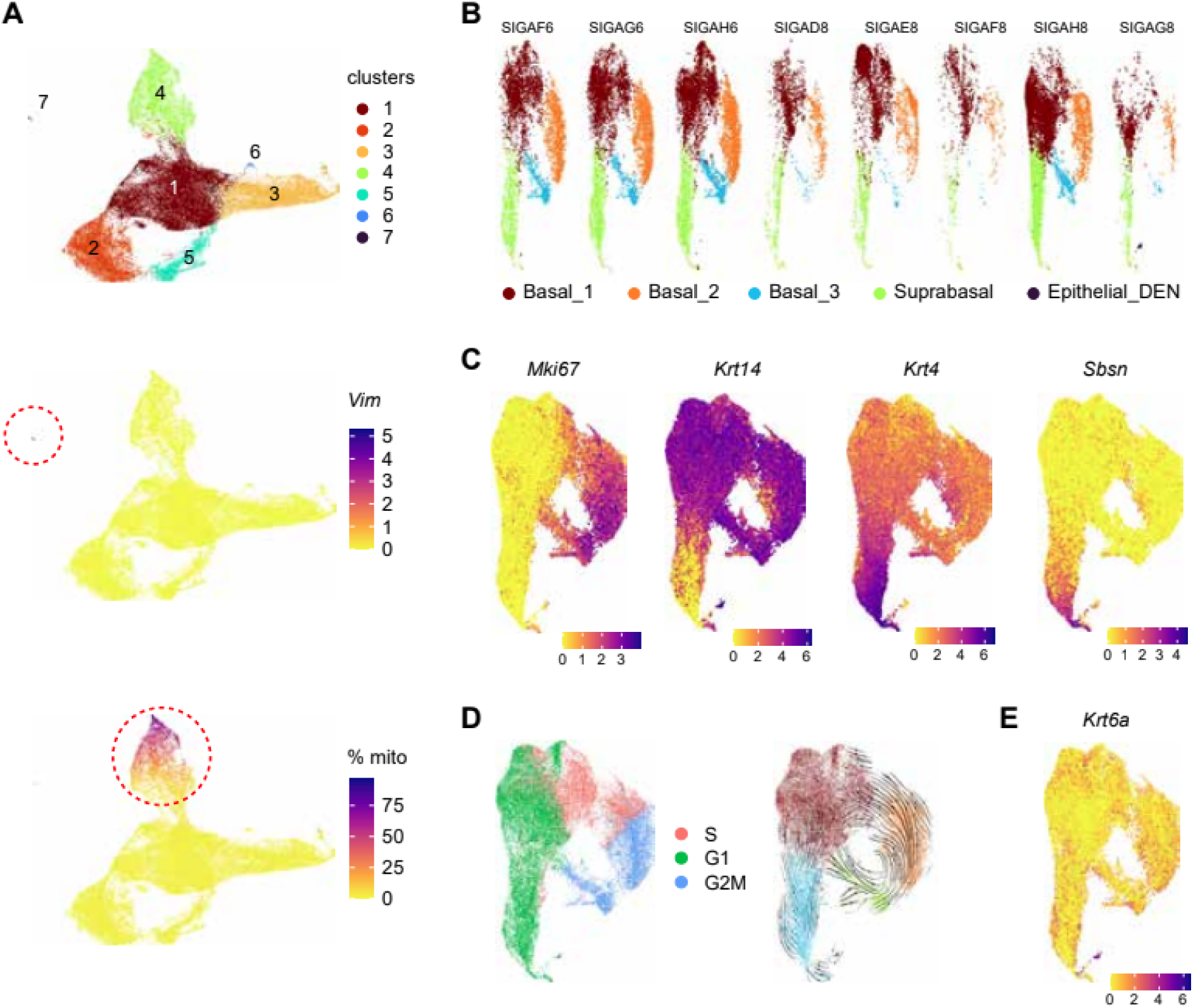
Characterization of single-cell transcriptomes in mouse esophagus. (A) UMAP embedding showing main clusters of individual cell transcriptomes before removal of mesenchymal cells (cluster 7) and likely damaged epithelial cells (cluster 4) (resolution = 0.2). Normalized Vimentin (*Vim*) gene expression values and % mitochondrial gene content per cell are highlighted in second and third panels, respectively. (B) UMAP embeddings of the subset of epithelial single-cell transcriptomes after re-normalization, separated per individual libraries. (C) Normalized gene expression values of some epithelial canonical markers: proliferative (*Mki67*), basal (*Krt14*), suprabasal (*Krt4*, *Sbsn*). (D) Cell-cycle phase inference (left panel) and RNA velocity stream vectors (right panel) obtained with scVelo. (E) Normalized *Krt6a* gene expression.

**Suppl. Fig. 2.**
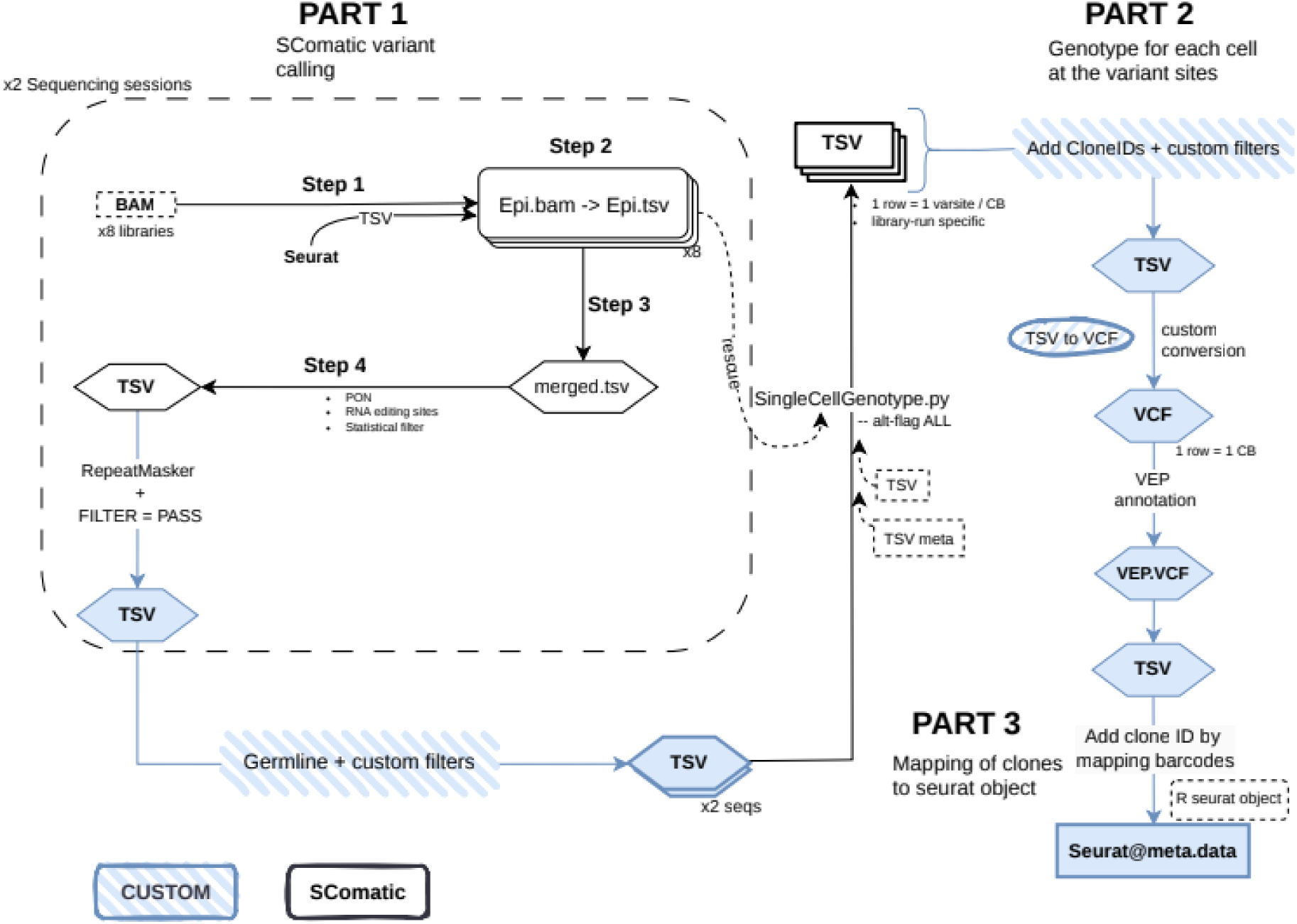
Flowchart of the analysis for variant detection, cellular mapping and functional annotation from mouse esophageal scRNA-seq data. BAM files from individual libraries in each sequencing session restrained to epithelial populations were taken for SComatic variant calling without further splitting (part 1). Variant calls underwent custom filters and were further processed to compute cellular genotypes and map clones to the seurat object (parts 2 and 3). SComatic inbuilt mainstream steps are shown in black, other steps colored.

**Suppl. Fig. 3.**
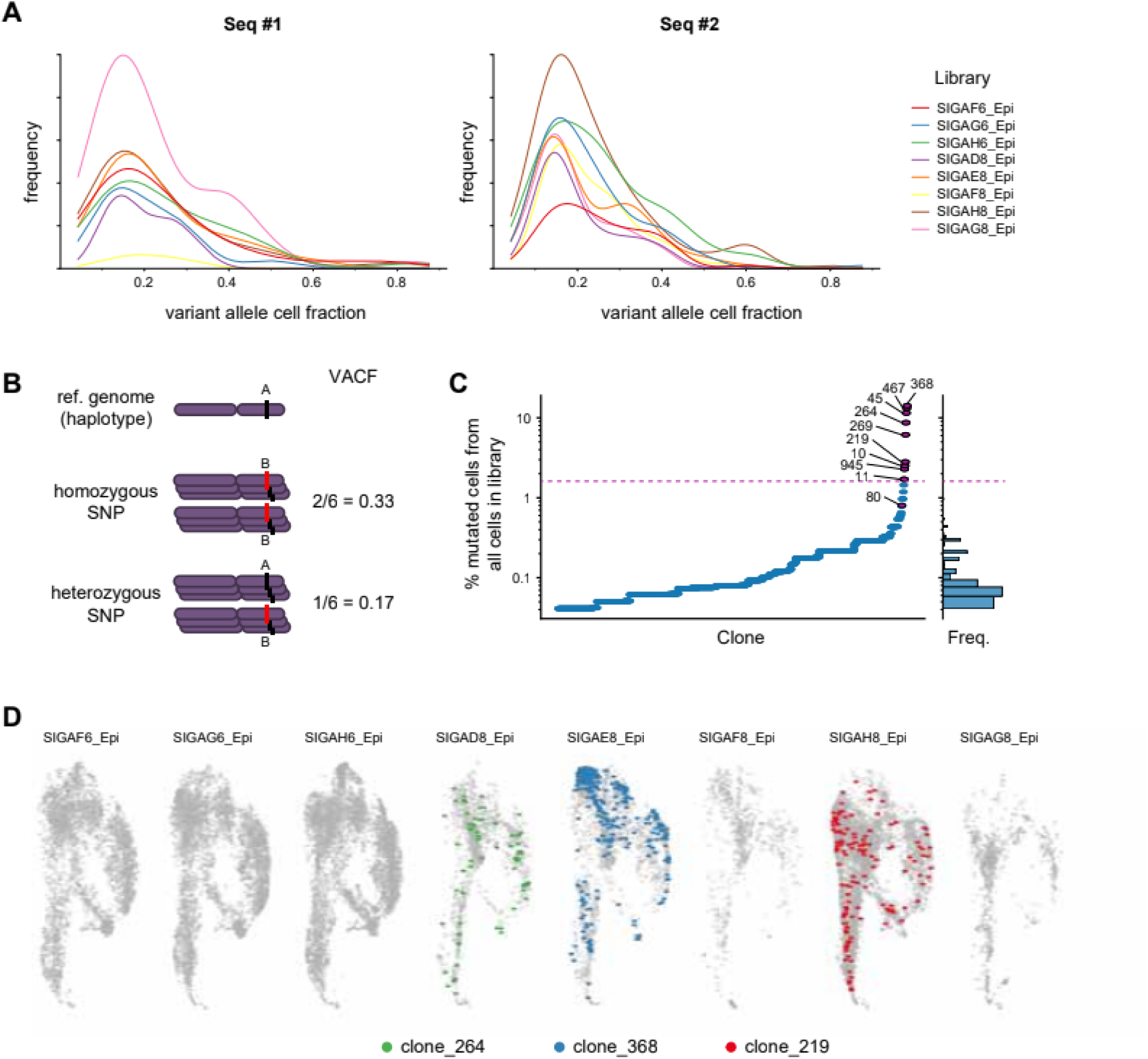
Filtering of repetitive-site calls and suspicious germline variants in mouse scRNA-seq data. (A) Distribution of the average fraction of mutated cells relative to the total number of cells with detectable on-site read coverage per library (*VACF*; SComatic *CCF* parameter), for variants found across multiple libraries in either sequencing session #1 or #2, respectively. (B) Scheme illustrating the fraction of mutated cells (*VACF*) expected for a widespread, animal-specific heterozygous or homozygous polymorphism with very limited read coverage, where just one allele is captured per individual cell (average on-site read depth right above 1). Notice individual libraries come from 3 mice each. (C) Fraction of mutant cells relative to total library size (total number of cells) for the different library-specific filtered variants, with mutant clones sorted in ascending size order. Points shown in red (above the threshold) correspond to outlier clones that are removed as suspicious germline variants (R getOutliers package with default settings) (clone ID numbers displayed). (D) Cellular mapping of some suspicious germline variants onto UMAP embeddings. Cells with alternative base calls for the given site (clone IDs) are colored.

**Suppl. Fig. 4.**
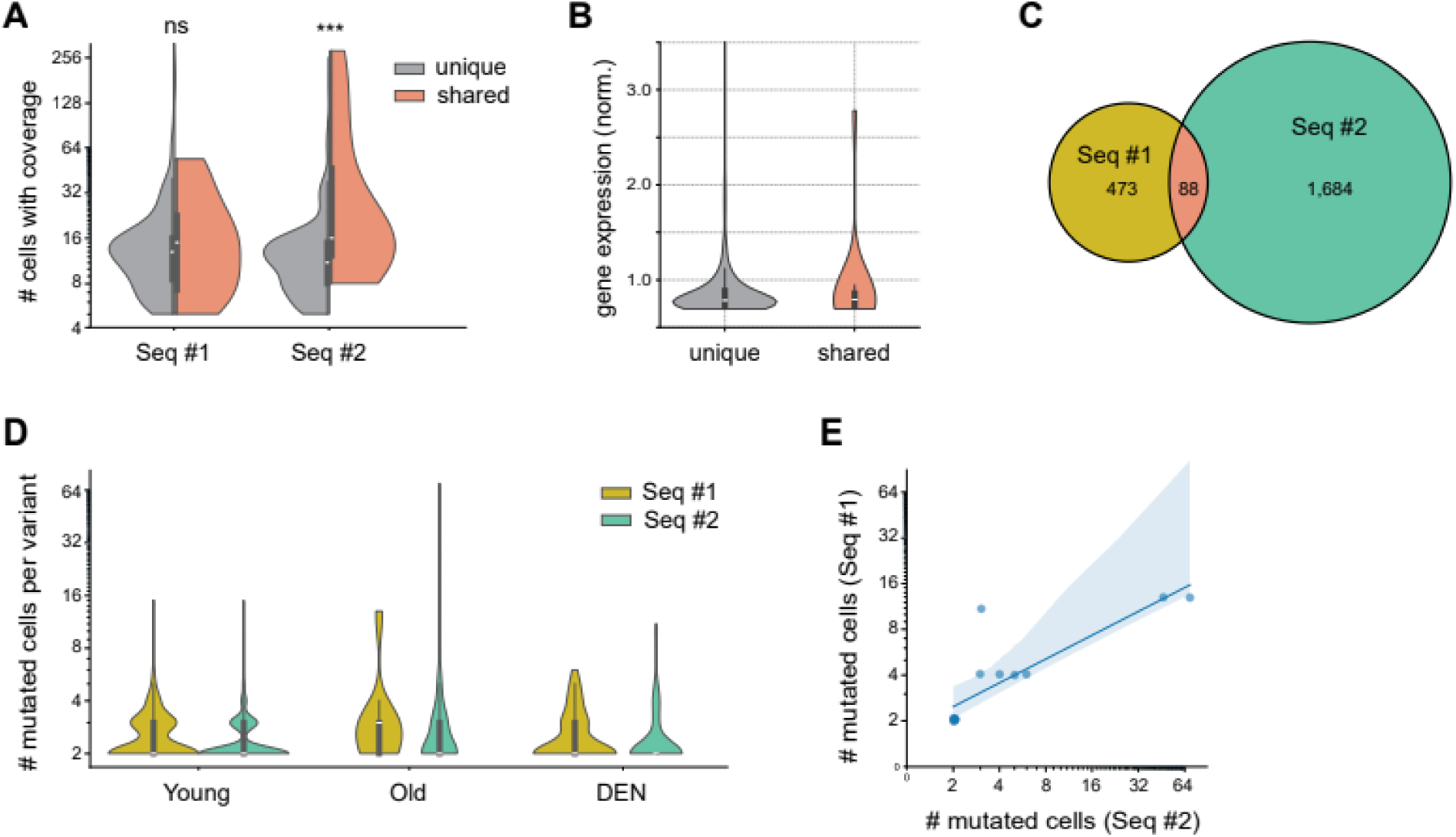
Exploratory analysis of candidate somatic mutations in mouse. (A) Number of cells with detectable on-site read coverage (SComatic *Nc* parameter), and (B) normalized gene expression values across cells, for variants found mutated in a single session versus those in both sessions. Cellular statistics are divided by sequencing session (*p*-val: 0.249, <0.001 for seq #1 and #2, respectively; one-sided Mann-Whitney *U*-test). (C) Venn diagram illustrating the number of unique and shared mutated cell barcodes detected across sequencing runs. (D) Distribution of apparent clone sizes (number of mutant cells) for variants found in the different experimental conditions. Data divided by sequencing session. (E) Comparison of apparent clone sizes (number of mutant cells) found for shared mutations in one versus another sequencing session (Pearson correlation: *R*=0.96). Notice higher apparent sizes in sequencing session #2 are consistent with its higher sequencing depth.

**Suppl. Fig. 5.**
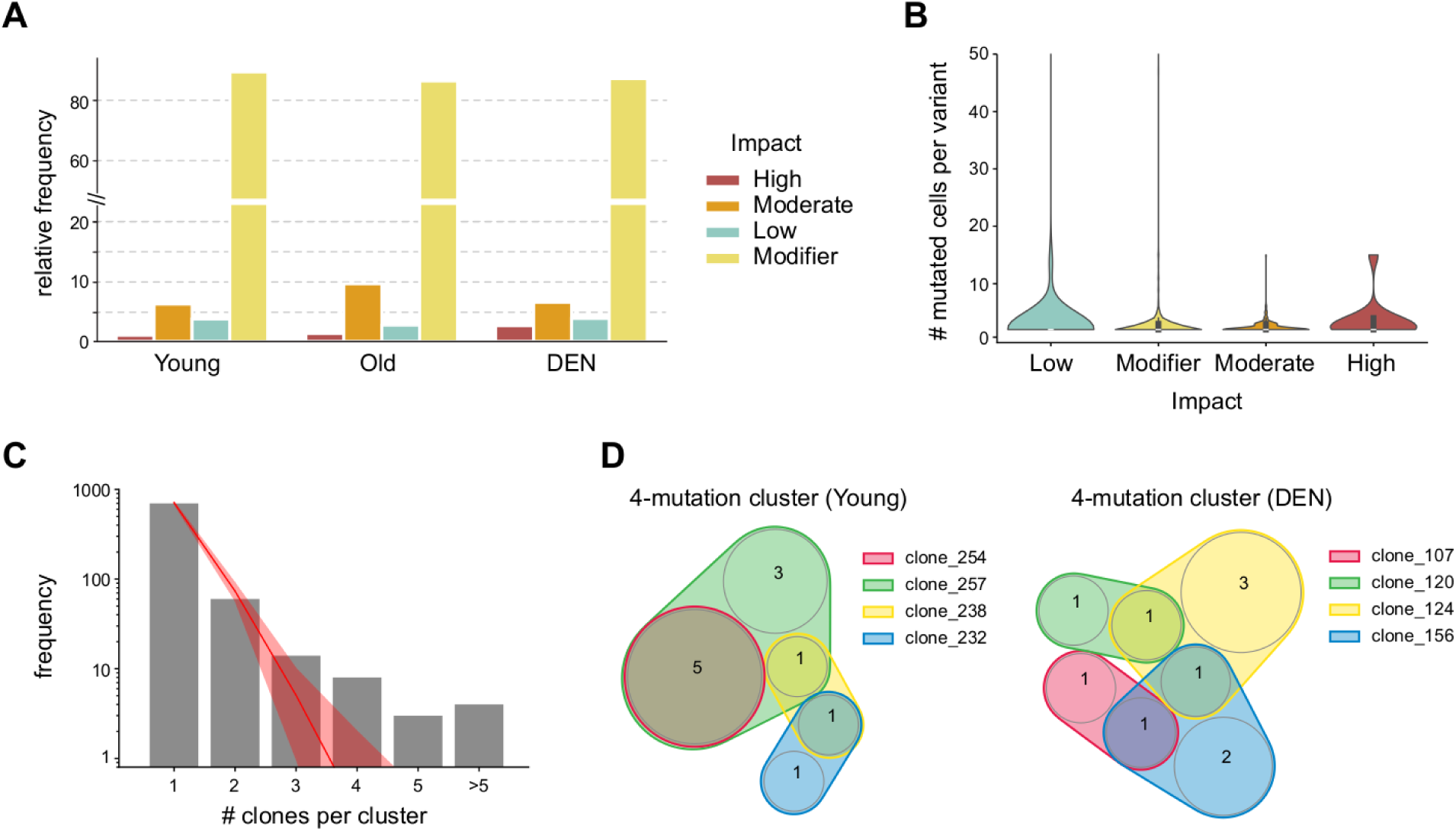
Functional impact and clonal structure of mutant cell populations in mouse. (A) Relative frequency of different levels of impact across mutations in the different experimental conditions, following VEP annotation (**Methods**). (No significant differences in DEN vs. control; Fisher exact test). (B) Distribution of apparent clone sizes (number of mutant cells) for variants annotated with each level of impact. (C) Frequency of individual mutation co-localization across cells, showing involvement in increasing size clusters. Cluster sizes are given by the number of individual mutations sharing at least one overlapping mutant cell in common (**Methods**). Best-fit Poisson distribution (expected rate: λ=0.207) is shown overlaid, with 95% uncertainty margins given by the actual sampling error (bootstrapping). (D) Venn diagrams of two examples of complex mutant clusters, illustrating partial cellular overlap between constituent individual mutations. Numbers of common and exclusive cells are displayed on the diagram.

**Suppl. Fig. 6.**
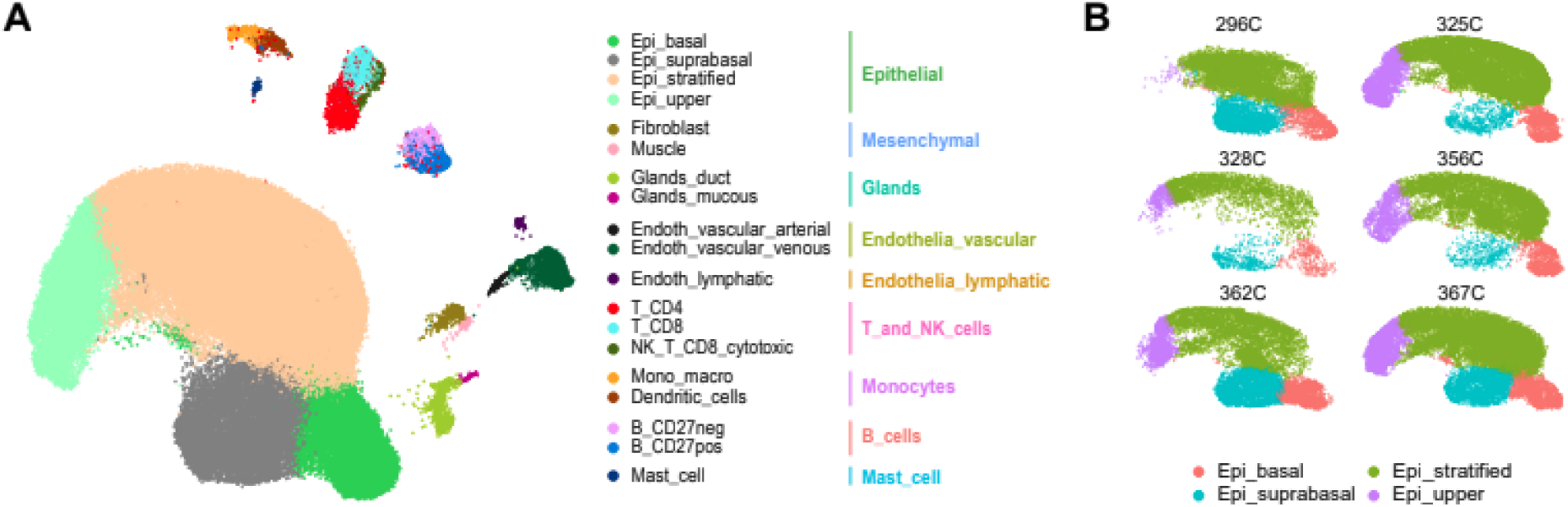
Annotation of single-cell transcriptomes in human esophagus. (A) UMAP-embedding of high-quality cell transcriptomes in the dataset, highlighting cell-type annotations used in the original publication from Madissoon et al (2019) and their correspondence with the more summarized ones used in our study (right-most legend). (B) UMAP-embedding subsetting epithelial cells from each donor, highlighting distribution of the 4 different epithelial subtypes.

**Suppl. Fig. 7.**
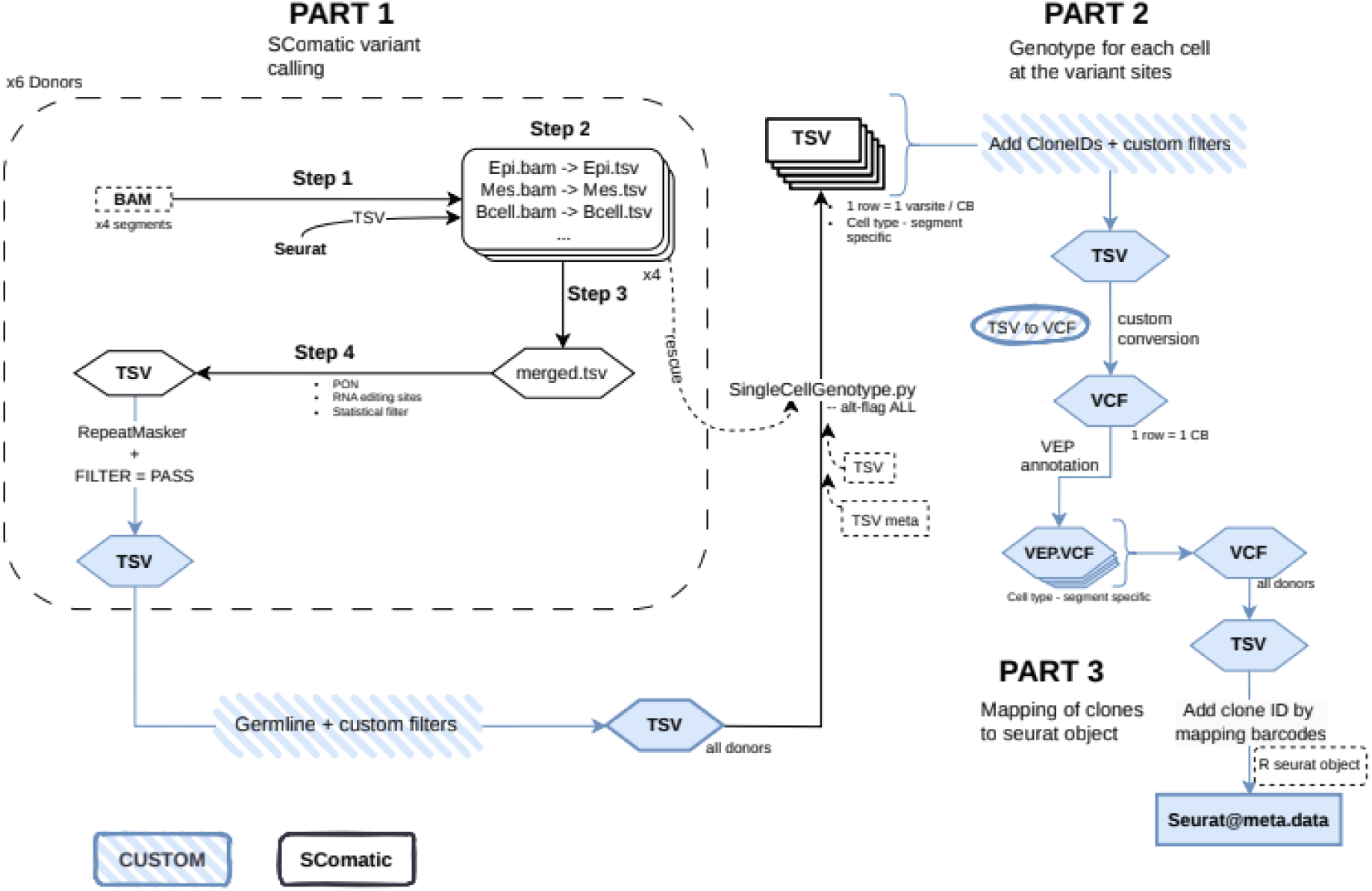
Flowchart of the analysis for variant detection, cellular mapping and functional annotation from human esophageal scRNA-seq data. BAM files from each individual donor were splitted into cell type- and segment-specific BAM files for SComatic variant calling (part 1). Variant calls underwent custom filters and were further processed to compute cellular genotypes and map clones to the seurat object (parts 2 and 3). SComatic inbuilt mainstream steps are shown in black, other steps colored.

**Suppl. Fig. 8.**
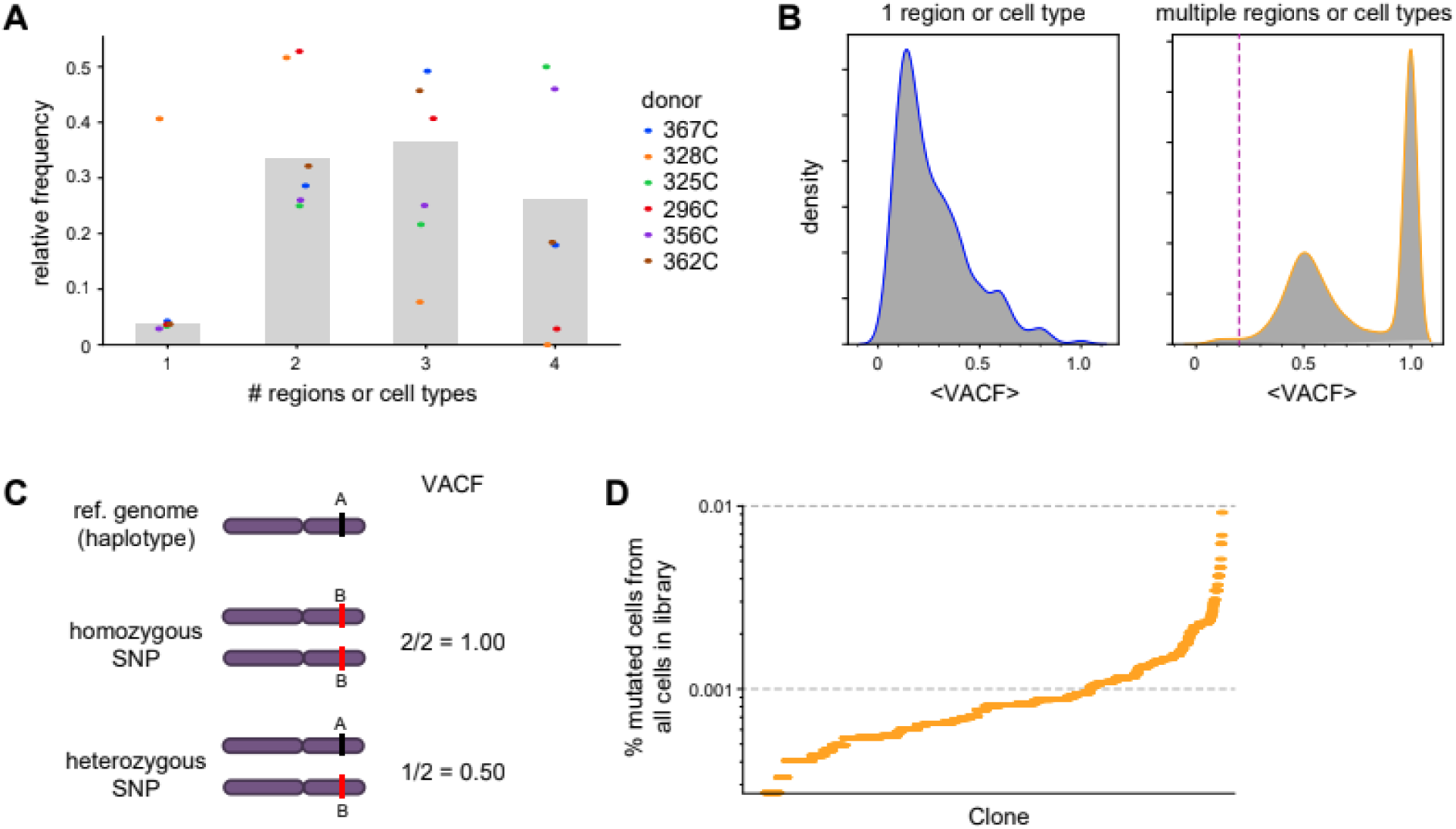
Filtering of suspicious germline variants in human scRNA-seq data. (A) Proportion of non-repetitive site variants found in 1, 2, 3 or 4 cell type-segments (bars). Dots correspond to relative frequencies separated per individual donor. (B) Distribution of the average fraction of mutated cells relative to the total number of cells with detectable on-site read coverage per library (*VACF*; SComatic *CCF* parameter) for variants supported by a single (left panel) or multiple (right panel) cell type-segments. The *VACF*=0.2 cutoff value applied to restrict to somatic mutations among the 2 cell type-segment variants is shown. (C) Scheme illustrating the fraction of mutated cells (*VACF*) expected for a widespread, individual-specific heterozygous or homozygous polymorphism with very limited read coverage, where just one allele is captured per individual cell (average on-site read depth right above 1). Notice individual libraries are donor specific. (D) Fraction of mutant cells relative to total library size (cells) for the different filtered variants, with mutant clones sorted in ascending size order. Values remain homogeneously low with no large outliers.

**Suppl. Fig. 9.**
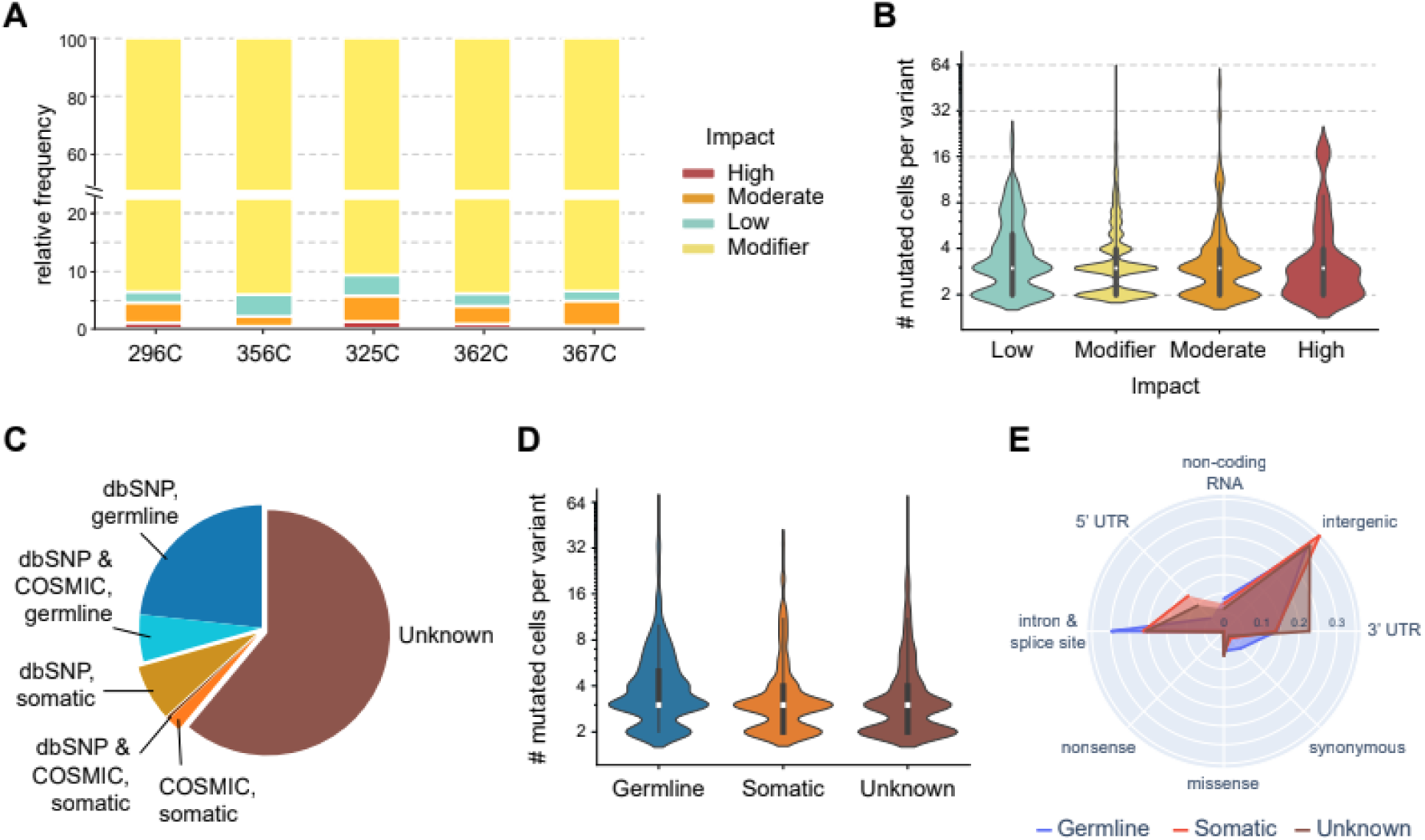
VEP annotation and exploratory analysis of candidate somatic mutations in human. (A) Relative frequency of different levels of impact across mutations in the different individual donors, following VEP annotation (**Methods**). (B) Distribution of apparent clone sizes (number of mutant cells) for variants annotated with each level of impact. (C) Classification of candidate mutations based on existing evidence in variation databases (**Methods**). (D) Distribution of apparent clone sizes (number of mutant cells) for variants classified as germline, somatic and unknown (*p*-val: 2.2e-5, 1.6e-12, 0.34 for germline vs somatic, germline vs unknown, somatic vs unknown, respectively; one-sided Mann-Whitney *U*-test). (E) Scatter-polar histogram showing relative frequencies of variants mapped to different gene subregions/consequence types, separated for the three distinct classes of mutations (*p*-val: 1.3e-4, 2.0e-14, 0.05 for germline vs somatic, germline vs unknown, somatic vs unknown, respectively; Pearson Chi-square test, with B-H correction for multiple testing).

**Supp. Fig. 10.**
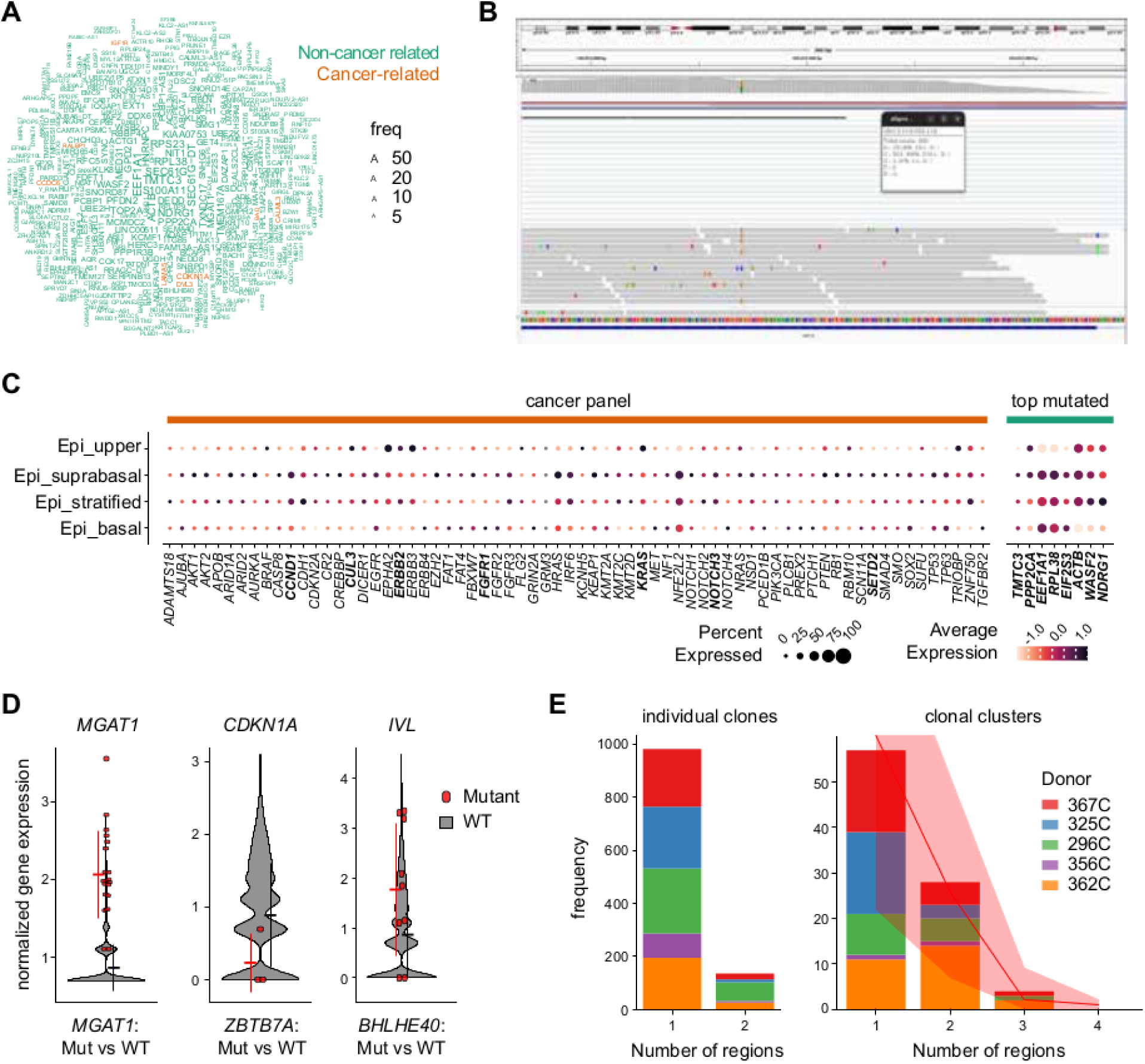
Limitations of mutation detectability on cancer-related genes and clonal structure features inferred from human scRNA-seq. (A) Word-cloud of top 400 most frequently mutated genes from human scRNA-seq. Names are scaled by incidence (total number of cells with variant calls for that gene). Cancer-associated genes are highlighted in orange. (B) Screenshot from Integrative Genomics Viewer (IGV) illustrating an example of a scRNA-seq derived variant mapped to 3’ UTR of highly-expressed *RFC5* gene on chr12_118032119 position (HGVSc: ENST00000454402.7:c.*841C>A). (C) Average normalized gene expression and detectability across epithelial single-cell transcriptomes of cancer-associated genes listed in Martincorena et al (2018) panel and top mutated genes from our analysis. Dot size indicates the % of cells where the gene was detected. (D) Normalized expression for some genes potentially affected by detected mutations: *MGAT1* in response to a *cis*-acting modifier mutation onto its 5’ UTR region (HGVSc: ENST00000307826.5:c.-222C>G) (left panel); *CDKN1A* in response to a moderate-impact, missense mutation onto its repressor, *ZBTB7A* transcription factor (HGVSc: ENST00000322357.9:c.1382C>T) (mid panel); *IVL* in response to a high-impact, stop-gained mutation onto *BHLHE40* transcription factor controlling keratinocyte differentiation (HGVSc: ENST00000256495.4:c.382+1G>A) (right panel). Normalized gene expression values in individual mutant cells (in red) are compared with those of non-mutant cells from the same subepithelial clusters (grey). Errorbars stand for mean ± SD (Only cells with non-null expression values of the mutated gene are considered) (*p*-val: 1.03E-19, 0.109, 0.027, respectively; two-sided Mann-Whitney *U*-test). (E) Relative frequencies of individual mutant clones (left panel) and clone clusters (right panel) spanning one or multiple territories or biopsies. Expected frequency values assuming random clone association appear overlaid in red (permutation test by bootstrapping).

## SUPPLEMENTARY TABLES

**Table S1. Variant calling file (VCF) of fully annotated mutations in mouse esophageal scRNA-seq data.**

**Table S2. Gene-set enrichment analysis output with top enriched pathways related with scRNA-seq-derived mutations in mouse.**

**Table S3. Annotation of clones mutant for cancer-associated genes in mouse.**

**Table S4. Variant calling file (VCF) of fully annotated mutations in human esophageal scRNA-seq data.**

**Table S5. Gene-set enrichment analysis output with top enriched pathways related with scRNA-seq-derived mutations in human classified as germline vs. somatic and unknown.**

**Table S6. Metadata relating file names to sample identification codes and features in the human dataset.**

